# Overexpression of NIMA-related kinase suppresses cell proliferation and tip growth in a liverwort *Marchantia polymorpha*

**DOI:** 10.1101/2023.01.25.525476

**Authors:** Hikari Mase, Yoshihiro Yoshitake, Takayuki Kohchi, Taku Takahashi, Hiroyasu Motose

## Abstract

NIMA-related kinases (NEKs) regulate a series of mitotic events in fungi and animals, whereas plant NEKs regulate growth direction of cells and organs. The liverwort *Marchantia polymorpha* has a single functional Mp*NEK1* gene, whose knockout leads to twisted growth of rhizoids. Mp*NEK1* is also expressed in the meristem of vegetative flat organ, thallus, while its function remains unknown. Here, we generated transgenic lines for the inducible expression of Mp*NEK1* using an estrogen receptor mediated system. Estradiol treatment efficiently induced the accumulation of Mp*NEK1* mRNA and MpNEK1-Citrine fusion protein throughout plant body. Overexpression of MpNEK1 severely suppressed growth of rhizoids and thalli, eventually causing the lethality of juvenile plants. The effect of estradiol was reversible until 3 days, whereas 7-days treatment resulted in irreversible suppression of growth. This severe effect was observed even at the nanomolar level of estradiol. EdU staining and microtubule imaging clearly indicated the suppression of cell proliferation by estradiol-induced MpNEK1. Unexpectedly, the overexpression of kinase-deficient MpNEK1 also suppressed thallus growth and rhizoid formation, despite their slightly mild effect than the full length MpNEK1, indicating phosphorylation-independent mechanism of growth suppression. In conclusion, overexpression of MpNEK1 suppresses cell division and elongation, leading to growth cessation and lethality. Our results imply that the expression of MpNEK1 is tightly regulated and plant NEKs might control cell division as in fungi and animals.

## Introduction

Never in mitosis A (NIMA)-related kinase (NEK) is a Ser/Thr protein kinase that controls a series of mitotic events in fungi and animals (O’Connell et al. 2003, Fry et al. 2012), while plant NEK members have been shown to regulate directional growth (Motose, et al. 2008; Sakai, et al. 2008). *Arabidopsis* NEK6 suppresses ectopic outgrowth of epidermal cells through microtubule organization and interaction with other NEK members (Motose, et al. 2011). NEK6 localizes to the shrinking ends of microtubules, phosphorylates five amino acid residues of β-tubulin, and depolymerizes aberrant cortical microtubules to promote epidermal cell elongation (Takatani, et al. 2017). Furthermore, NEK6 has been shown to stabilize growth direction of hypocotyls through selectively removing cortical microtubules that best align with maximal tensile stress (Takatani, et al. 2020). NEK6-mediated attenuation of tensile-stress response increases the noise in microtubule orientation to buffer mechanical conflicts and to stabilize growth direction.

The liverwort *Marchantia polymorpha* is an emerging model to study evolution of land plants (Kohchi, et al. 2021) because of its low genetic redundancy (Bowman, et al. 2017) and various genetic tools (Ishizaki, et al. 2015; Ishizaki, et al. 2016). Indeed, *Arabidopsis* has seven NEK members, whereas *M. polymorpha* has a single *NEK* gene, Mp*NEK1* (Takatani, et al. 2015; Otani, et al. 2018). Thanks to these advantages, MpNEK1 has been shown to direct tip growth of rhizoids through microtubule depolymerization in the apical region (Otani, et al. 2018). Rhizoids are filamentous rooting cells, mainly elongating from the ventral epidermis and are thought to be required for plant anchorage to the soil and water/nutrient uptake (Jones and Dolan 2012; Shimamura, 2016; Kohchi, et al. 2021; Kanda, et al. 2022). Mp*nek1* mutant rhizoids frequently change their growth direction, resulting in the zigzag morphology (Otani, et al. 2018) and the defect of invasive growth to enter the hard substratum (Mase, et al. 2022). MpNEK1 localization in the apical microtubule foci and the increased microtubule stability in Mp*nek1* mutants demonstrate that MpNEK1 reorganizes and destabilizes apical microtubules to maintain growth polarity of rhizoids (Otani, et al. 2018). Similar phenotype was observed in the mutants of a microtubule-associated protein, WAVE DANPENED-LIKE (MpWDL) (Honkanen, et al. 2016; Champion, et al. 2021). MpWDL might stabilize longitudinal microtubules in the shank region of rhizoids. Thus, microtubule organization and dynamics are essential for directional growth of rhizoids.

However, it is not clear how MpNEK1 regulates microtubules to stabilize growth direction. Furthermore, functional significance of Mp*NEK1* expression in the meristem of thalli remains unclear. The gain of function approach could be helpful to elucidate these problems and to identify downstream proteins that are phosphorylated by MpNEK1. Here, we adopted estradiol-inducible system to overdrive Mp*NEK1* and to estimate its effects on growth and development in *M. polymorpha*.

## Results

### Establishment of MpNEK1 inducible lines

In this study, we generated Mp*NEK1* inducible transgenic lines employing the XVE chimeric transcriptional activator (Zuo, et al. 2000). This activator is composed of the DNA-binding domain of LexA, the transcriptional activation domain of VP16, and the carboxyl estradiol binding domain of the human estrogen receptor. This system has no apparent toxic physiological effects and has been effectively used in flowering plants. The XVE vector was optimized for the application to *M. polymorpha* in several points. Because the endogenous elongation factor Mp*EF1α* promoter is more active in the meristem and sexual organ than the cauliflower mosaic virus *35S* promoter (Althoff, et al. 2014), the Mp*EF1α* promoter was utilized to drive XVE. However, only a few research has been reported on the application of XVE system in *M. polymorpha* (e.g. Flores-Sandoval, et al. 2016; Furuya, et al. 2022).

We isolated eight transgenic lines by transformation of wild type sporelings with pMpGWB344-MpNEK1 (MpEF1α pro:XVE>>LexAop:MpNEK1), in which XVE is expressed under the control of Mp*EF1α* promoter and activates transcription of Mp*NEK1* through the binding to the LexA operator in the presence of estradiol. Among them, seven transgenic lines (line #2 to 8) were most identical to the wild type in the growth and morphology of thalli. These transgenic lines were examined for the efficiency of Mp*NEK1* induction by quantitative real-time PCR (RT-qPCR) (Supplementary Fig. S1). Two-week-old thalli of the wild type (WT) and transgenic lines were transferred to the medium with or without β-estradiol and incubated for 24 hours. The expression levels of Mp*NEK1* were remarkably increased in most transgenic lines (#2, 4, 5, 7, and 8). Mp*NEK1* expression was 4-15 fold higher in estradiol treatment than in the estradiol-free medium. Estradiol did not induce Mp*NEK1* expression in the wild type and line #6. Mp*NEK1* induction was well reproducible and reliable, especially in line #2, 4, and 8, which exhibit no obvious defects in growth and morphology without estradiol. Thus, these were used for the further analyses.

### MpNEK1 induction severely suppresses thallus growth

We conducted phenotypic analysis to access the effects of estradiol-induced MpNEK1 on growth and development. The gemmae of the wild type and Mp*NEK1* inducible lines were grown in the presence or absence of exogenous β-estradiol for 24 days. There was no obvious effect of β-estradiol on the growth and morphology of the wild type thalli (Supplementary Fig. S2). In the inducible lines, thallus growth was severely suppressed when cultured with β-estradiol (Fig. 1). Most gemmalings stopped growing and died in the presence of estradiol (Fig. 1), whereas the line #6 did not show growth retardation (Supplementary Fig. S3). It was noteworthy that Mp*NEK1* was not induced in the line #6 (Supplementary Fig. S1). These results clearly demonstrate that induction of Mp*NEK1* severely suppresses thallus growth and causes juvenile lethality.

**Fig 1.**
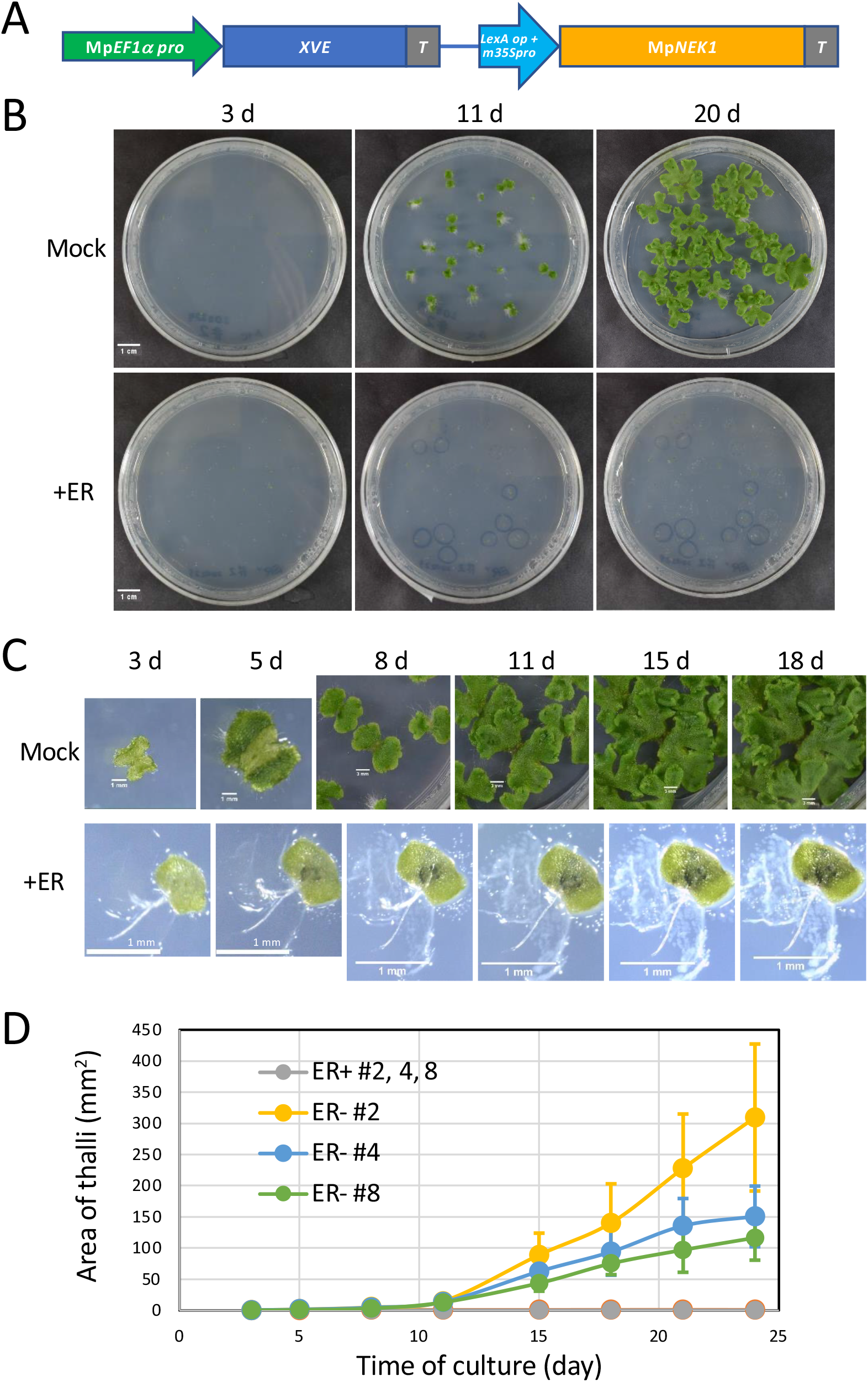
Effect of estradiol on thallus growth of Mp*NEK1* inducible line. (A) Construct for Mp*NEK1* induction. Mp*EF1a* promoter, terminator (T), LexA operator (LexA op), minimal CaMV35S promoter (m35Spro). The diagram is not shown to scale. (B) The gemmae of Mp*NEK1* inducible line (#2) were planted in the agar medium with (+ER) or without 10 µM estradiol (Mock) and grown for 3, 11, and 20 days. (C) Time course of growth of Mp*NEK1* inducible line (#2) with (+ER) or without 10 µM estradiol (Mock) (D) Quantification of thallus growth of Mp*NEK1* inducible line (#2, 4, 8) with (+ER) or without 10 µM estradiol (Mock). The mean projection area of thalli (n = 8-11 plants) was quantified by ImageJ. Circles and error bars indicate mean values and standard deviations, respectively.

We next examined whether effect of Mp*NEK1* induction is reversible or not. The gemmalings of the wild type and Mp*NEK1* inducible lines were grown in the presence of estradiol, and then transferred to the estradiol-free medium at various times after estradiol treatment (Fig. 2). After 1 and 3-day treatment with estradiol, thallus growth seemed to be recovered, whereas irreversible growth suppression was observed after 7 and 14-day treatment. In the wild type, growth inhibitory effect of estradiol was not observed. Therefore, one-week Mp*NEK1* induction (estradiol treatment) confers irreversible effect on thallus growth.

**Fig 2.**
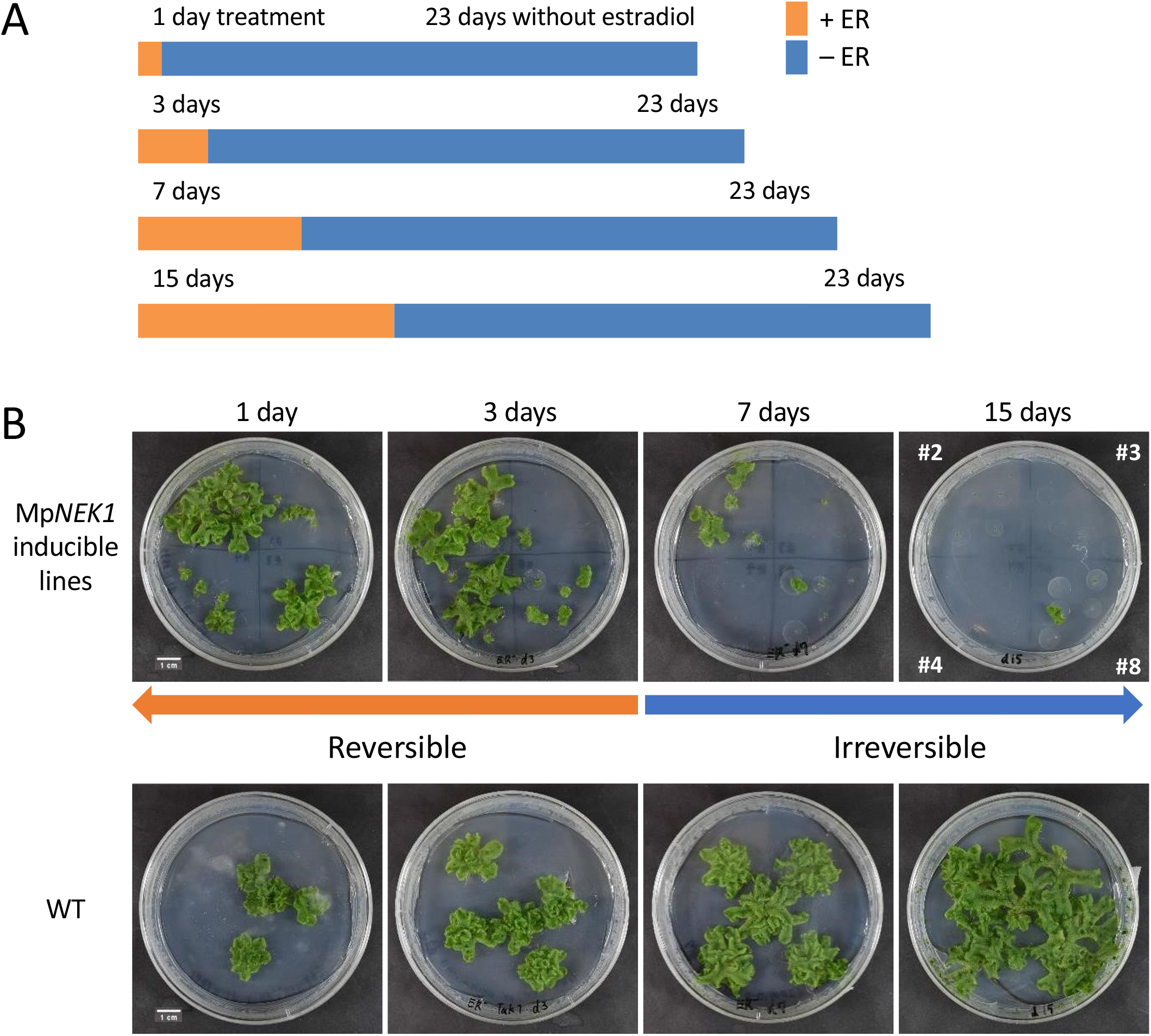
Reversible or irreversible effect of estradiol on thallus growth (A) Schematic diagram of treatments. Gemmae were planted in the agar medium with 10 µM estradiol, grown in the same medium for 1, 3, 7, or14 days, and then transferred to and grown in the estradiol-free medium for 23 days. (B) Gemmae of the wild type (WT) and four independent inducible lines (#2, 3, 4, and 8) were grown as shown in (A). The periods of estradiol treatment is shown above the photographs taken at the 23 day after transfer to the estradiol-free medium.

In the XVE induction system in flowering plants, 2 to 10 µM concentrations of estradiol have been commonly used. In the Mp*NEK1* inducible lines, 10 µM estradiol caused severe growth suppression and lethality. However, more mild conditions would be desirable to further investigate MpNEK1 function. To determine less-effective conditions and dose-dependent effect of estradiol, thalli of the wild type and Mp*NEK1* inducible lines were incubated in the presence of estradiol at various concentrations (Fig. 3). The wild-type thallus showed no obvious phenotype in all concentration ranges, whereas severe growth suppression was observed in the Mp*NEK1* inducible lines even at the 10 and 100 nM of estradiol. From this result, Mp*NEK1* induction has the potent effect on thallus growth.

**Fig 3.**
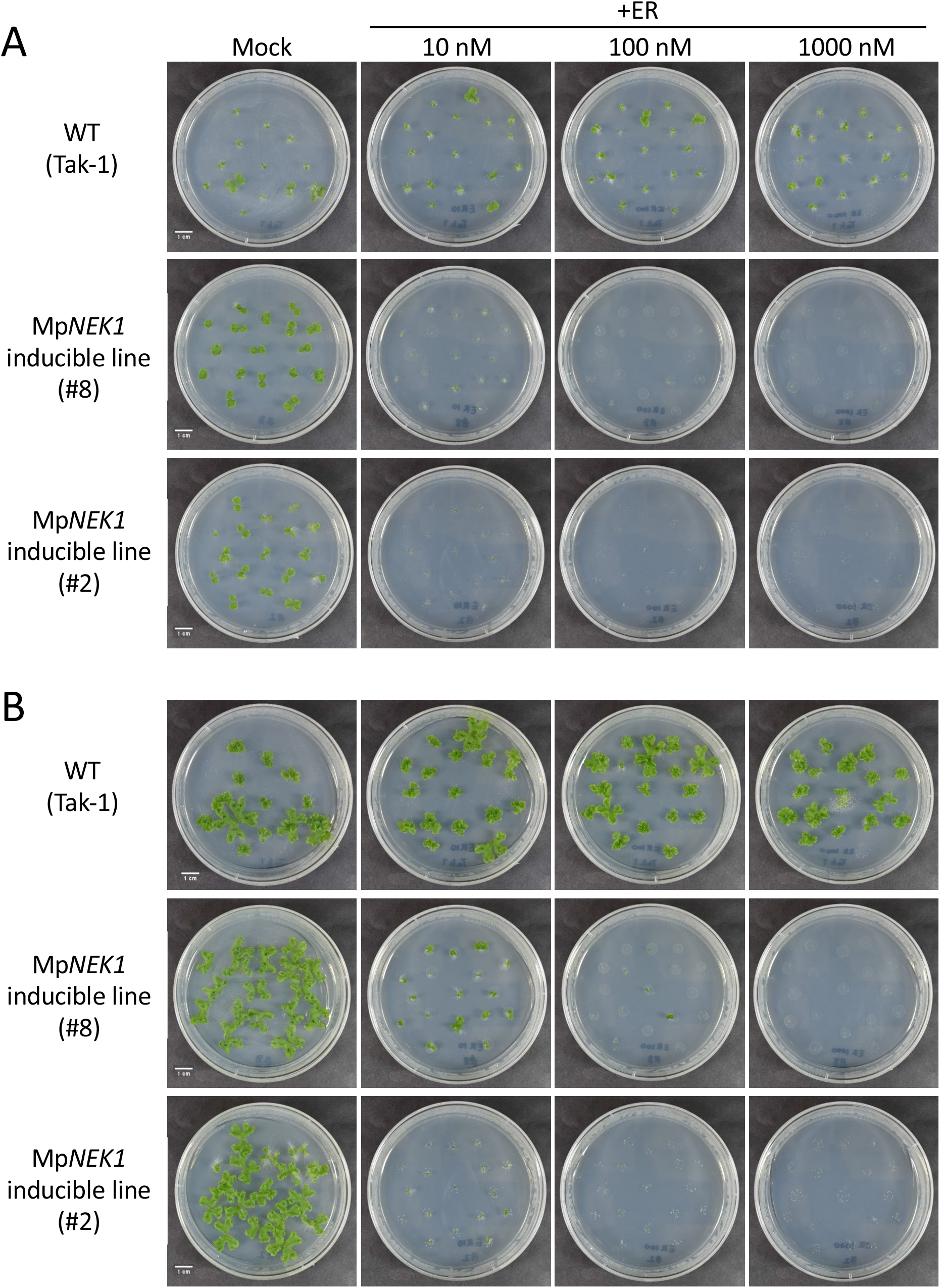
Effect of various concentrations of estradiol on thallus growth. The gemmae of the wild type and Mp*NEK1* inducible lines were planted in the agar medium supplemented with or without estradiol at the concentration of 10 nM, 100 nM, and 1000 nM and grown for 13 (A) and 16 days (B).

We noticed that some plants obtained the resistance to estradiol and survived in the agar medium supplemented with estradiol (Fig. 4). The estrogen resistance was observed in the next generation gemmae, suggesting cell-cell movement of siRNA from the thallus to gemma. These estradiol-resistant plants appeared more frequently in the lower concentrations and exhibited no induction of Mp*NEK1* (Supplementary Fig. S4). At the lower concentrations, larger numbers of thallus cells could survive and acquire estradiol resistance by gene silencing mechanism. Thus, lower concentrations of estradiol are not desirable conditions, which often induce resistant plants.

**Fig. 4.**
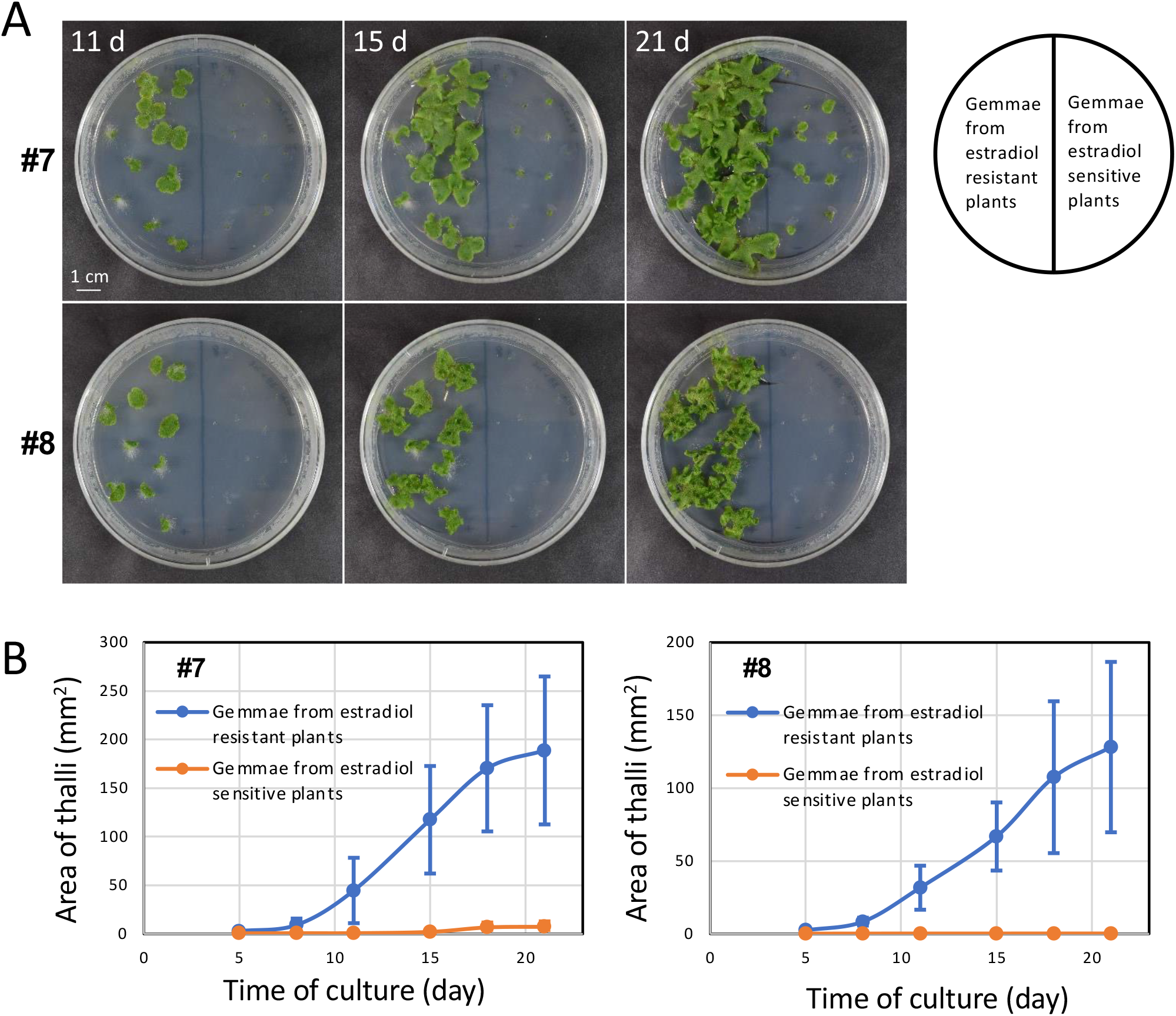
Estradiol resistance and its transfer to the next generation. (A) The gemmae of the estradiol resistant plants or sensitive plants of Mp*NEK1* inducible lines were planted in the agar medium supplemented with 10 µM estradiol and grown for 11, 15, and 21 days. (B) Quantification of growth of gemmalings derived from the estradiol resistant plants or sensitive plants of Mp*NEK1* inducible lines. The mean projection area of thalli (n = 10 plants) was quantified by ImageJ. Circles and error bars indicate mean values and standard deviations, respectively.

### MpNEK1 induction suppresses cell proliferation

Previous studies show that plant NEKs mainly regulate growth direction of cells and organs. However, severe suppression of thallus growth by estradiol-induced Mp*NEK1* suggests some roles of plant NEK in cell division and proliferation. To determine whether estradiol-induced Mp*NEK1* affects cell division, we monitored cell proliferation activity by staining with EdU (5-ethynyl-2’-deoxyuridine) (Fig. 5). This thymidine analog can be incorporated into DNA during S phase. The subsequent reaction of EdU with a fluorescent azide labels newly replicated DNA strands of the cells in S phase during the incubation with EdU.

**Fig. 5.**
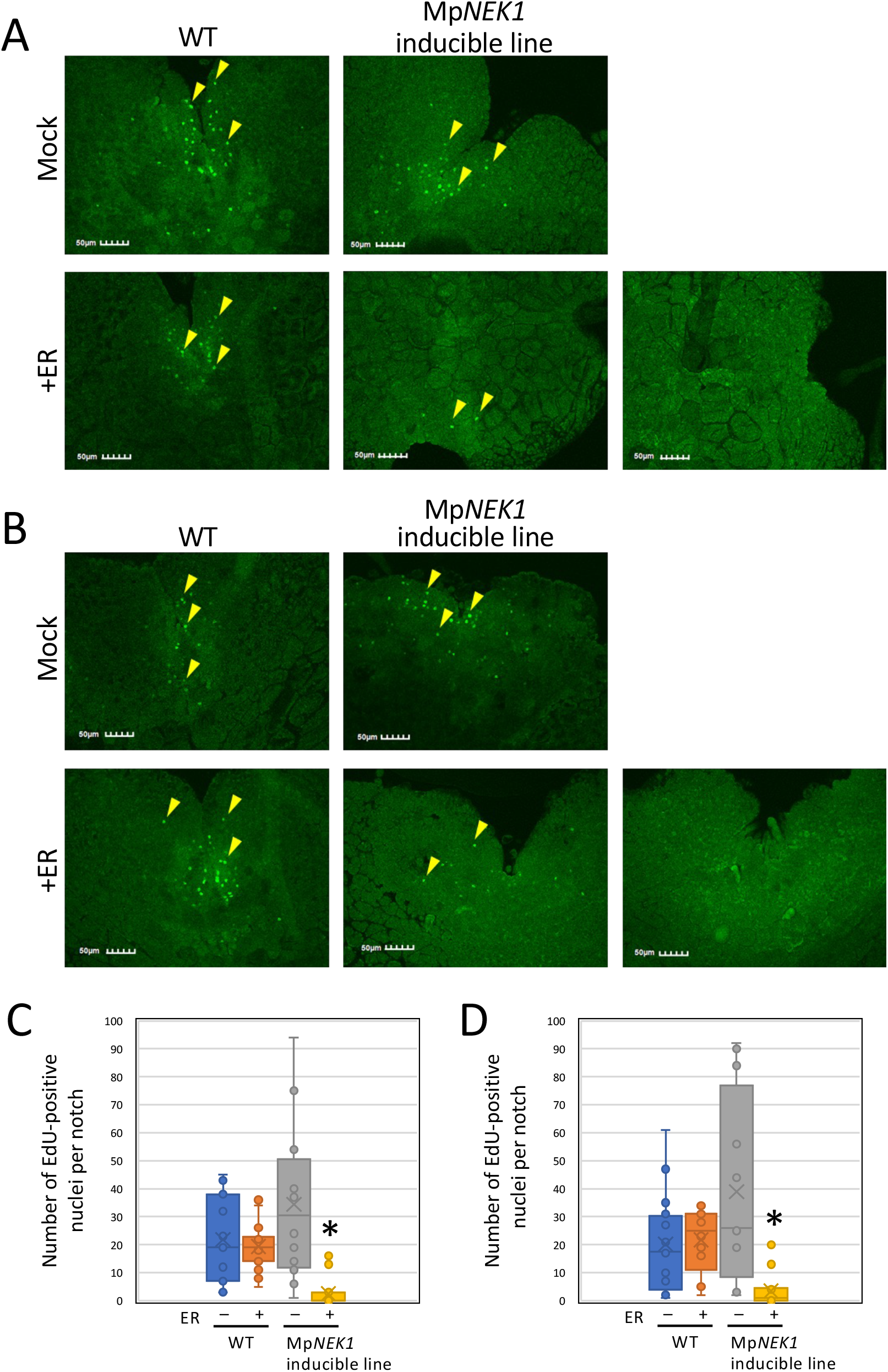
Effect of estradiol on cell proliferation (A) EdU-labelled nuclei in the gemmalings treated with 1 µM estradiol for 4 days. The gemmae of the wild type (WT) and Mp*NEK1* inducible line were planted in the medium supplemented with or without 1 µM estradiol, grown for 4 days, and incubated with 10 µM EdU for 1 hour in liquid medium of the same kinds. EdU-labelled nuclei were visualized according to the manufacture’s instruction as described in methods. (B) EdU-labelled nuclei in the gemmalings treated with 1 µM estradiol for 1 day. The gemmae of the wild type (WT) and Mp*NEK1* inducible line were planted in the medium without estradiol, grown for 3 days, and then transferred to and grown in the medium supplemented with or without 1 µM estradiol for 1 day. The gemmalings were incubated with 10 µM EdU for 1 hour in liquid medium of the same kinds. EdU-labelled nuclei were visualized according to the manufacture’s instruction as described in methods. (C) Quantification of EdU-labeled nuclei in the thalli grown as in (A). Data are shown by box plots (n = 12-16 plants). An asterisk indicates significant difference from the control (-ER) (*t*-test, P < 0.05). (D) Quantification of EdU-labeled nuclei in the thalli grown as in (B). Data are shown by box plots (n = 12-16 plants). An asterisk indicates significant difference from the control (-ER) (*t*-test, P < 0.05).

In both the wild type and inducible lines, EdU-positive nuclei were observed in the meristematic notch region in thalli grown in the absence of estradiol (Fig. 5). The amount of EdU-positive nuclei was almost similar in the wild type with or without estradiol, whereas it was significantly reduced in the estradiol-treated inducible lines. This result demonstrates that estradiol-induced Mp*NEK1* suppresses cell proliferation.

### MpNEK1 induction suppresses mitotic cell division

Because *Arabidopsis* NEK6 depolymerizes specific microtubules, which detached from plasma membrane or aligned to the maximal tensile stress, to regulate cell expansion and organ growth (Takatani, et al. 2017; Takatani, et al. 2020), we monitored the effect of MpNEK1 overexpression on microtubule organization and mitotic apparatus. MpNEK1 inducible constructs were introduced into the transgenic lines expressing a microtubule marker Citrine-MpTUB2. We obtained the MpNEK1 inducible lines expressing Citrine-MpTUB2. The gemmae of these transgenic plants were grown for 3 days in the estradiol-free medium and then applied with the liquid medium with or without 10 µM estradiol. After 24 and 48 hours of treatment, thallus cells were observed with confocal microscopy. In the estradiol-treated thalli, number of mitotic cells was significantly decreased compared with the mock-treated thalli (Fig. 6A, B). We classified mitotic cells according to mitotic structures, prospindle, spindle, and phragmoplast, since these structures are clearly distinguished under confocal microscopy (Buschmann et al. 2016). Despite the slight increase of cells with prospindle, there are no considerable differences in the ratio of mitotic structures (Fig. 6C), suggesting that MpNEK1 induction does not severely affect mitotic progression. We also found that cortical microtubules were not affected with or without estradiol (Fig. 6A). Because we could not detect microtubule signal in rhizoids of the obtained transgenic lines, it could not be determined whether MpNEK1 induction affects microtubule organization in rhizoids. In summary, overexpression of MpNEK1 suppresses mitotic cell division.

**Fig. 6.**
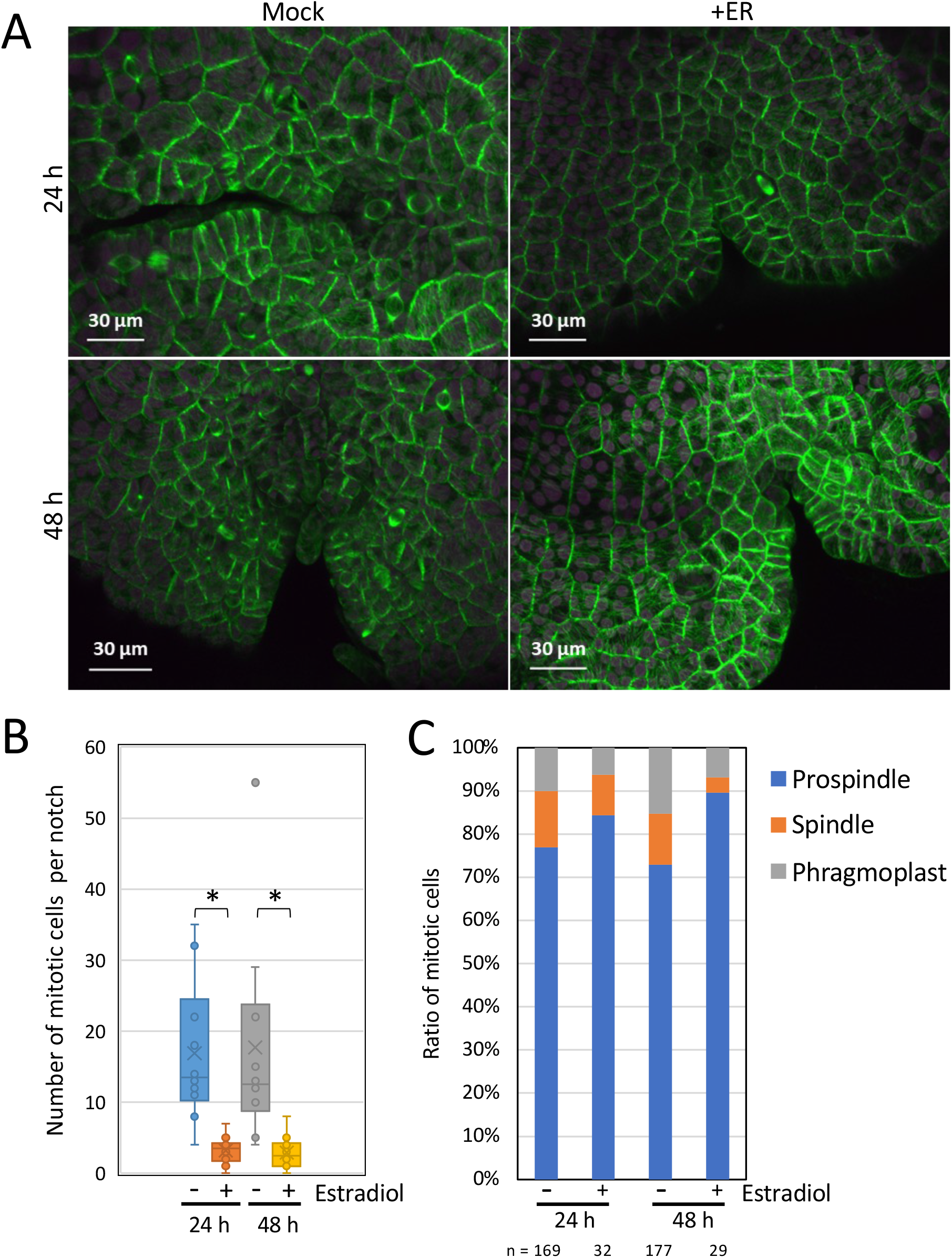
Effect of MpNEK1 induction on microtubules (A) The gemmae of MpNEK1 inducible line with a microtubule marker CaMV35S:Citrine-MpTUB2 were grown in the estradiol-free medium for 3 days and then applied with the liquid medium with (+ER) or without 10 µM estradiol (Mock). Thalli were observed under a confocal microscope after 24 h and 48 h of treatment. Green: Microtubule, magenta: plastid autofluorescence. (B) Number of mitotic cells per notch in the MpNEK1 inducible line with CaMV35S:Citrine-MpTUB2 after 24 h and 48 h of treatment with or without estradiol. Data are shown in the box plot (n = 10 notches, N > 6 plants). An asterisk indicates significant difference (t-test, P < 0.01). (C) The classification of mitotic cells with prospindle, spindle, or phragmoplast (n indicates the number of mitotic cells used for the classification).

### The overaccumulation of MpNEK1-Citrine suppresses growth of thalli and rhizoids

We generated inducible lines of MpNEK1-Citrine fusion protein (Fig. 7), in which protein induction and localization could be easily detected by confocal microscopy, facilitating functional analysis of MpNEK1. Eight transgenic lines were isolated by transformation of wild type sporelings with pMpGWB144-MpNEK1-Citrine. Among them, four transgenic lines (line #1, 5, 7, and 8) were most identical to the wild type in the absence of estradiol. When grown in the agar medium supplemented with estradiol, these transgenic lines exhibited severe growth retardation and juvenile lethality (Supplementary Fig. S5).

**Fig. 7.**
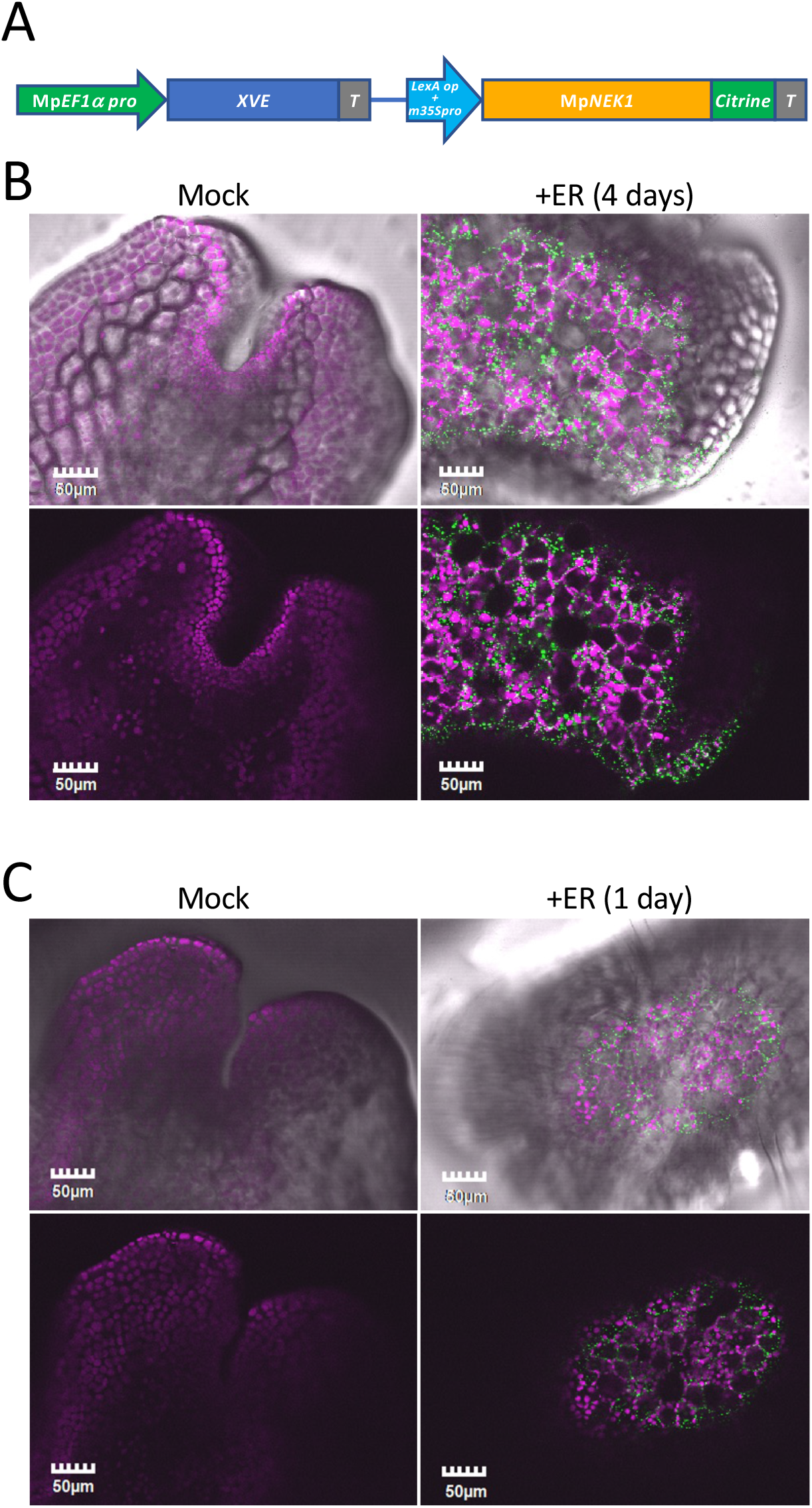
Localization of estradiol-induced MpNEK1-Citrine in the thallus (A) Construct for Mp*NEK1* induction. Mp*EF1a* promoter, terminator (T), LexA operator (LexA op), minimal CaMV35S promoter (m35Spro). The diagram is not shown to scale. (B) The gemmae of the Mp*NEK1* inducible line were planted in the medium with (+ER) or without 1 µM estradiol (Mock), grown for 4 days, and observed under a confocal microscope. (C) The gemmae of the Mp*NEK1* inducible line were planted in the medium without estradiol, grown for 2 days, and then added with (+ER) or without 1 µM estradiol (Mock) for 1 day. Upper panels are light field images merged with confocal images shown in the below panels. Green: MpNEK1-Citrine, magenta: plastid autofluorescence.

We next analyzed expression and localization of MpNEK1-Citrine fusion protein (Fig. 7B, C). The gemmae of MpNEK1-Citrine inducible lines were grown for 4 days in the medium with or without 1 µM β-estradiol and observed under a confocal microscope (Fig. 7B). MpNEK1-Citrine was not detected in the absence of estradiol, whereas it was accumulated throughout estradiol-treated thalli (Fig. 7B, Video S1). MpNEK1-Citrine localized in the cytoplasmic particles. In the 24-hour estradiol treatment (2-day old gemmalings grown without estradiol were treated with estradiol solution for 24 hours), similar expression and localization pattern were observed (Fig. 7C). From these results, MpNEK1-Citrine was induced throughout thalli within 24 hours after estradiol treatment and severely suppressed thallus growth.

We noticed accumulation of MpNEK1-Citrine particles in the meristem in some transgenic plants even in the absence of estradiol (Supplementary Fig. S6). This leaky expression levels were varied in each individual and transgenic line. In this study, we used plants and transgenic lines (line #1, 5, 7, and 8), which showed less leaky expression of MpNEK1-Citrine.

Taking accounts that MpNEK1 localizes in the apical microtubule foci to direct tip growth of rhizoids (Otani, et al. 2018), we analyzed effect and localization of estradiol-induced MpNEK1-Citrine in tip-growing rhizoids (Fig. 8). Because rhizoids were not formed when gemmae were grown with estradiol at the start of culture, we added estradiol solution on the 3-day-old gemmalings grown without estradiol to examine localization and effect of MpNEK1-Citrine in the time course. After 24-hour treatment, MpNEK1-Citrine accumulated in the apical region of rhizoids (Fig. 8A, Video S2). The expression level of MpNEK1-Citrine was varied in each rhizoid. A few rhizoids showed highly accumulated MpNEK1-Citrine, which formed large droplets, moved to the apical direction, and fused with each other (Video S3). To determine when MpNEK1-Citrine starts to accumulate, we observed rhizoids at the early periods of estradiol treatment (Fig. 8B). MpNEK1-Citrine began to accumulate at about 1 hour of treatment and gradually increased in the rhizoid apex. After 48 and 120 hours, MpNEK1-Citrine accumulated throughout the entire region of rhizoids in the particle and filamentous patterns (Fig. 8C, Video S4-S7). Tip growth was ceased or severely reduced after 48 and 120 hours of estradiol treatment (Fig. 8D, Video S4-6). After 48 hours, movement of MpNEK1-Citrine particles was obvious (Fig. 9A, Video S4, S5). MpNEK1-Citrine particles moved toward the apex (anterograde) and basal region (retrograde) at the same velocity (Fig. 9B). These results suggest that overaccumulation of MpNEK1 suppresses rhizoid growth.

**Fig. 8.**
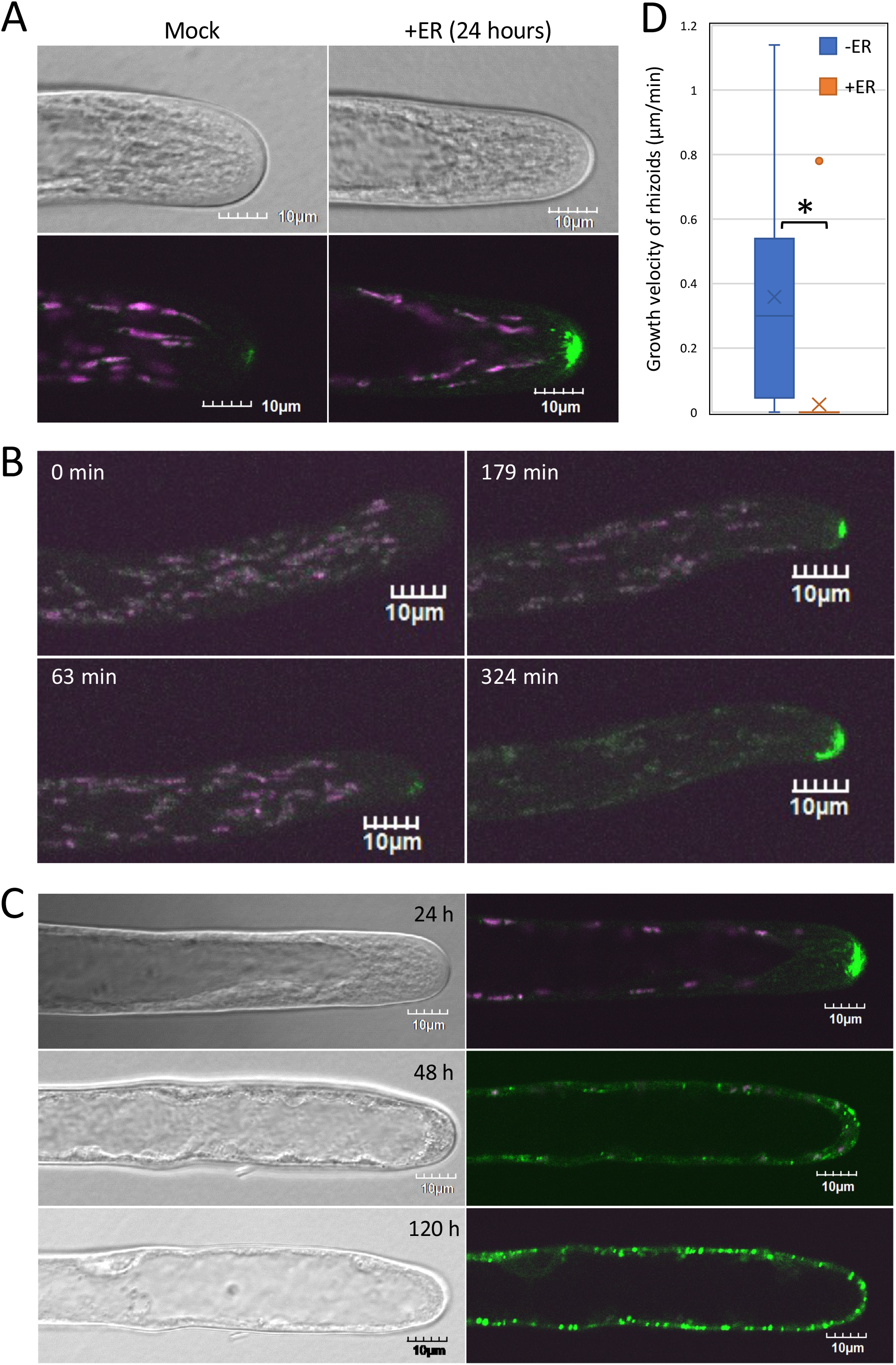
Localization of estradiol-induced MpNEK1-Citrine in rhizoids. (A) The gemmae of the Mp*NEK1* inducible line were planted in the medium without estradiol, grown for 3 days, and then applied with (+ER) or without 1 µM estradiol (Mock). Rhizoids were observed under a confocal microscope. Green: MpNEK1-Citrine, magenta: plastid autofluorescence. (B, C) The gemmae of the Mp*NEK1* inducible line were planted in the medium without estradiol, grown for 3 days, and then applied with 1 µM estradiol and observed at the time indicated in each panel. The same rhizoid was observed under a confocal microscope in the time course. Green: MpNEK1-Citrine, magenta: plastid autofluorescence. (D) Velocity of rhizoid growth in the MpNEK1-Citrine inducible line incubated for 48 hours with (+ER) or without 1 µM estradiol (-ER). Data are shown by box plots (n = 32 rhizoids). An asterisk indicates significant difference (*t*-test, P < 0.01).

**Fig. 9.**
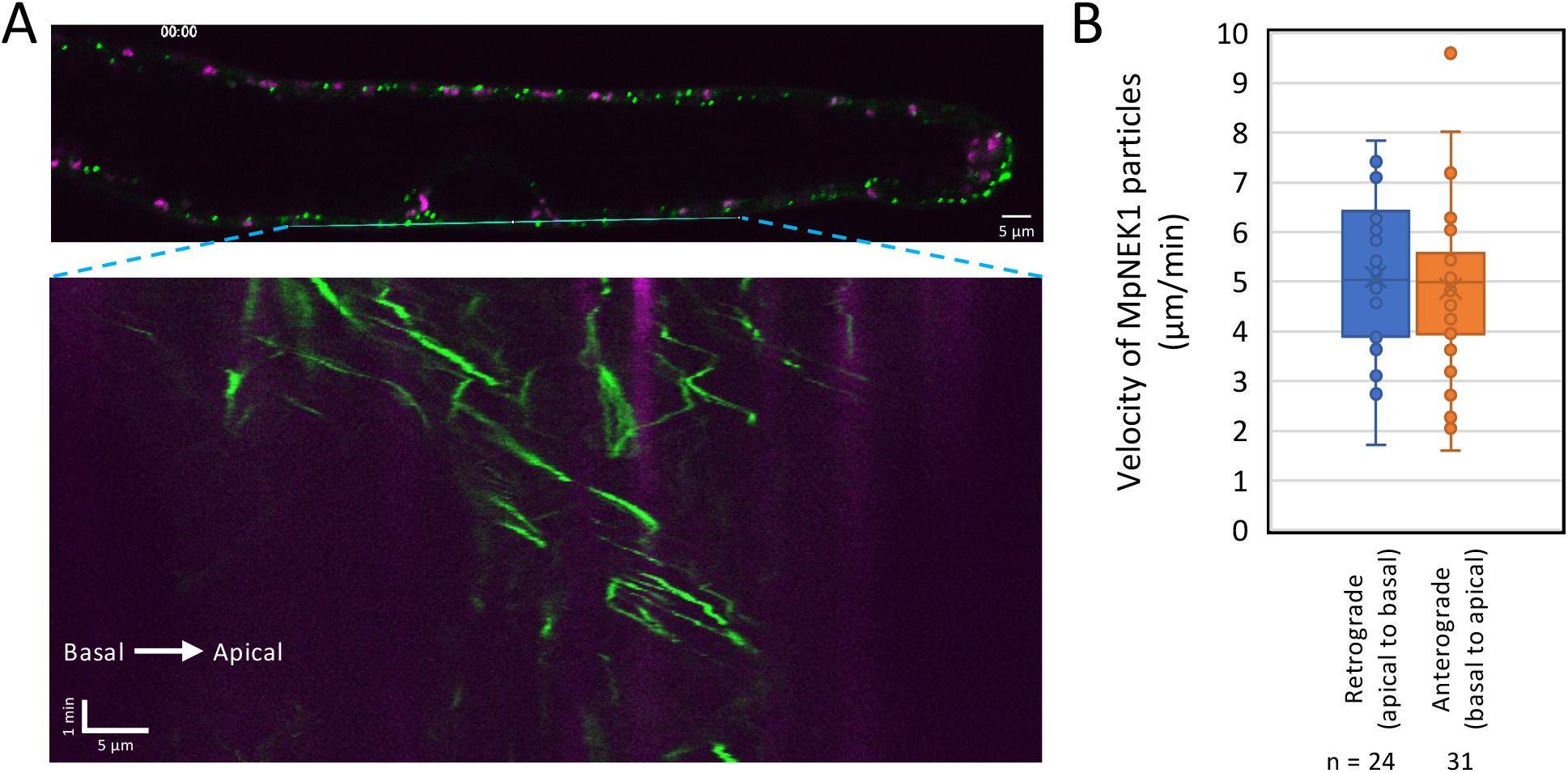
Movement of MpNEK1-Citrine particles in rhizoids. (A) The gemmae of the Mp*NEK1* inducible line were planted in the medium without estradiol, grown for 3 days, and then applied with 1 µM estradiol. Rhizoids were observed under a confocal microscope after 48 hours. The lower panel indicates kymograph of MpNEK1-Citrine particles. Green: MpNEK1-Citrine, magenta: plastid autofluorescence. (B) Velocity of MpNEK1-Citrine particles. Data are shown by box plots (n = 24 or 31 particles). There was no significant difference (*t*-test, P > 0.05).

### The overexpression of kinase-deficient MpNEK1 suppresses growth of thalli and rhizoids

To determine whether the kinase activity of MpNEK1 is required for growth suppression, we generated the inducible lines of MpNEK1 without its kinase activity (Fig. 10). For this purpose, lysine-34 essential for the kinase activity was substituted with glutamic acid (K37E, MpNEK1^K37E^) or the N-terminal kinase domain was deleted (KD deletion, MpNEK1^dKD^) (Fig. 10A). MpNEK1^K37E^ and MpNEK1^dKD^ were fused with Citrine at the C-termini and constructed into the XVE system as shown in Fig. 7A.

**Fig. 10.**
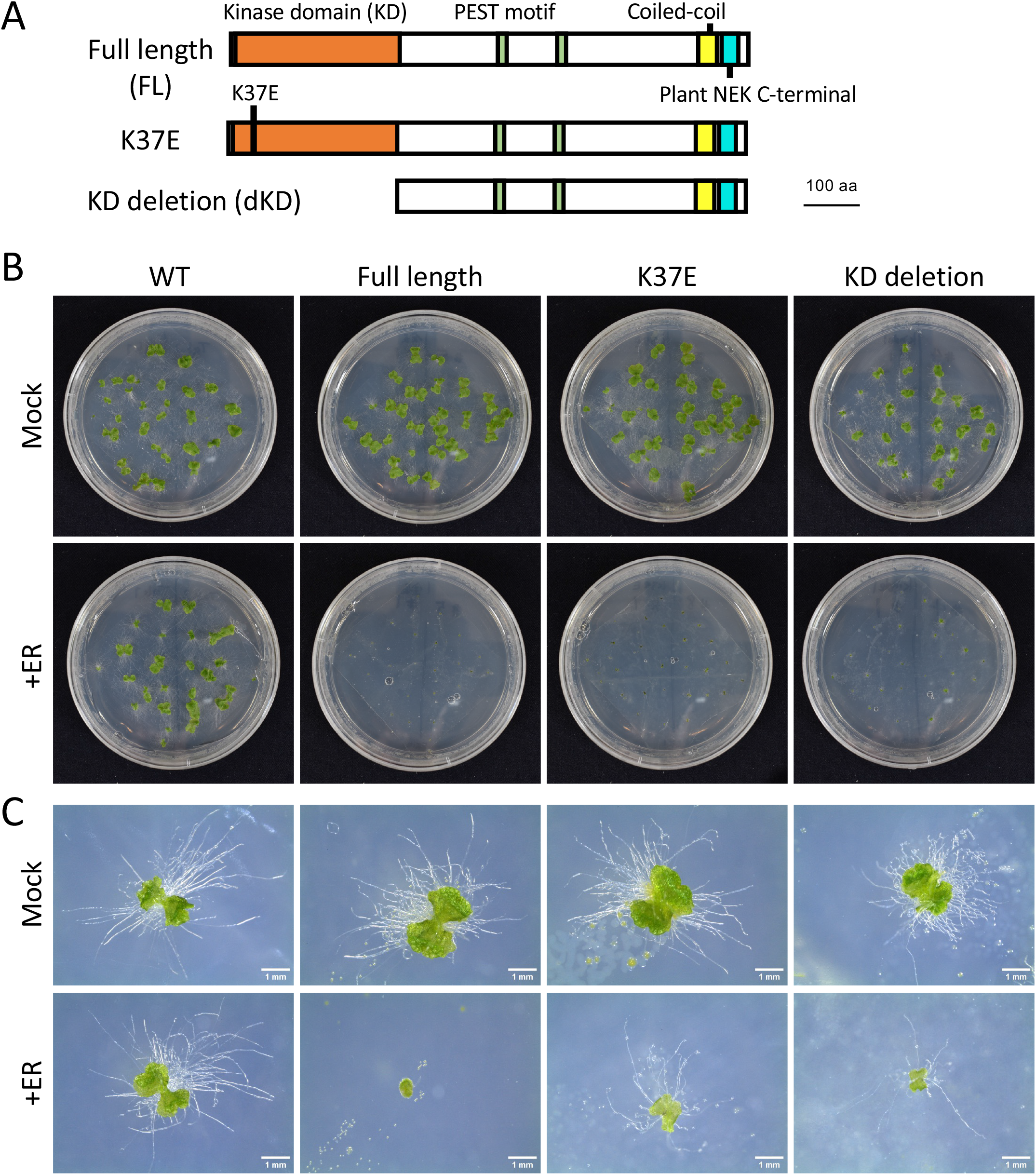
Effect of kinase-deficient MpNEK1 induction on the growth of thalli and rhizoids. (A) Full length and kinase-deficient MpNEK1 used for estradiol induction. (B) The gemmae of the wild type and inducible lines of full length MpNEK1, MpNEK1^K37E^, or MpNEK1^dKD^ (KD deletion) were planted in the medium with (+ER) or without 1 µM estradiol (Mock), and grown for 14 days. (C) The gemmae of the wild type and inducible lines of full length MpNEK1, MpNEK1^K37E^, or MpNEK1^dKD^ (KD deletion) were planted in the medium with (+ER) or without 1 µM estradiol (Mock), and grown for 7 days.

We compared the effects of MpNEK1^K37E^ and MpNEK1^dKD^ induction to that of the full-length MpNEK1 induction (Fig. 10, 11, Supplementary Fig. S7). The gemmae of the wild type and inducible lines of full length MpNEK1, MpNEK1^K37E^, and MpNEK1^dKD^ were grown in the medium with (+ER) or without 1 µM estradiol (Mock). In the presence of estradiol, thallus growth was suppressed in all inducible lines (Fig. 10B). This finding indicates that the kinase activity of MpNEK1 is not required for the suppression of thallus growth. However, close-up observation showed that the thalli of MpNEK1^K37E^ and MpNEK1^dKD^ inducible lines were larger than that of full-length MpNEK1 inducible line (Fig. 10C). Furthermore, rhizoids were scarcely formed in the full length inducible lines, whereas MpNEK1^K37E^ and MpNEK1^dKD^ inducible lines developed several rhizoids, which elongated as those of the wild type (Fig. 10C).

We quantified the number of rhizoids, which seems to be a good indicator of the difference among the inducible lines (Fig. 11). The gemmae of the wild type and inducible lines of full length MpNEK1, MpNEK1^K37E^, and MpNEK1^dKD^ were grown in the medium with (+ER) or without 1 µM estradiol (Mock) for 7 days (Fig. 11A). The number of rhizoids in the wild type was not affected by estradiol. Although the number of rhizoids was significantly decreased in all of the inducible lines, the inducible lines of MpNEK1^K37E^ and MpNEK1^dKD^ develop more rhizoids than the full length inducible lines. This difference was confirmed in the normalized data by using the average number of rhizoids in estradiol-free medium as a standard (Fig. 11B). The difference between kinase deficient and full length inducible lines was more clear in the quantification of rhizoids in 14-d treatment (Fig. 11C). Thus, the induction of kinase-deficient MpNEK1 is slightly milder than the full length induction. The kinase activity of MpNEK1 may have a little contribution to growth suppression.

**Fig. 11.**
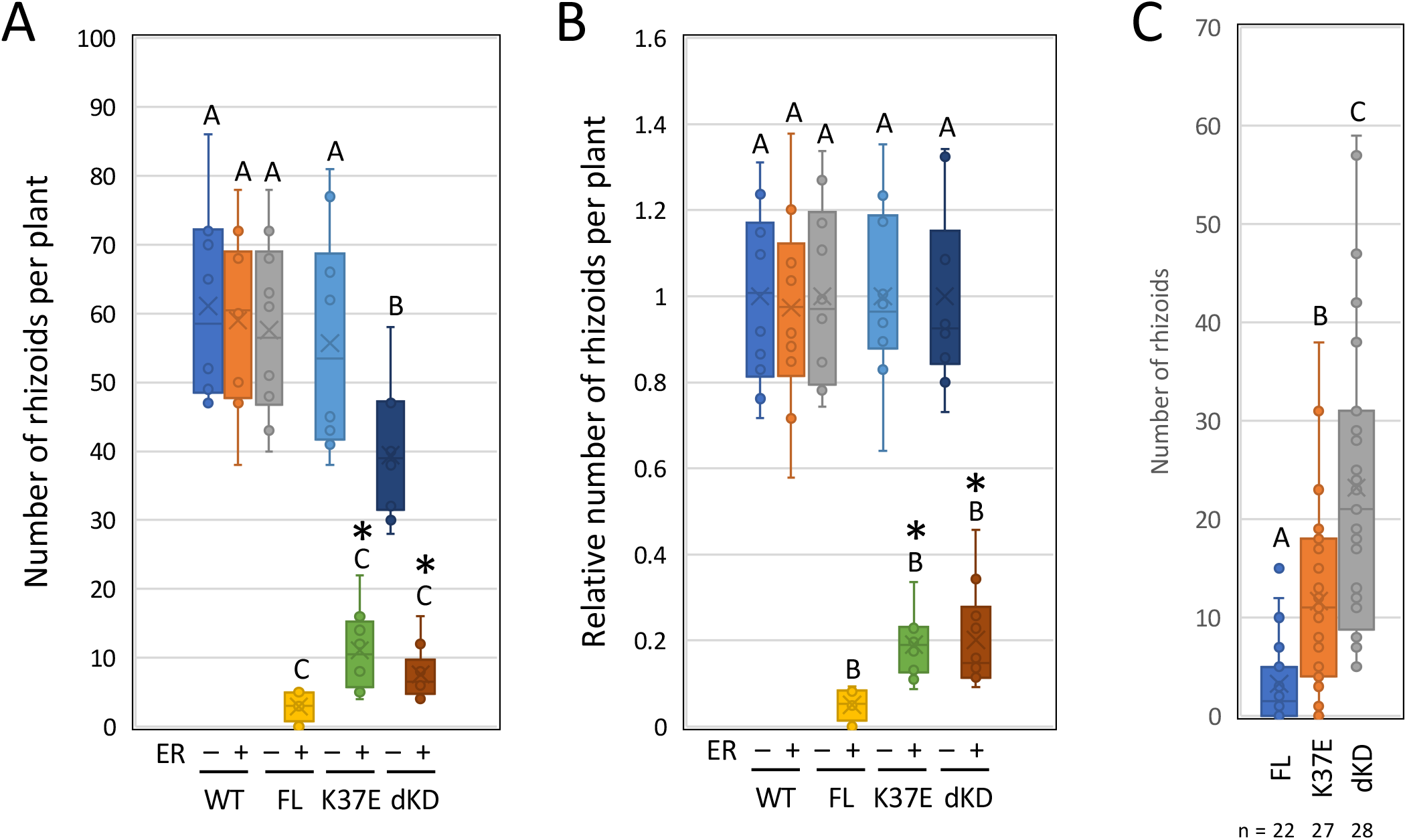
Effect of MpNEK1 induction on the number of rhizoids. (A) The number of rhizoids in the gemmae grown for 7 days in the medium with (+ER) or without 1 µM estradiol (Mock). The gemmae of the wild type (WT) and inducible lines of full length MpNEK1-Citrine (FL), MpNEK1^K37E^-Citrine (K37E), and MpNEK1^dKD^-Citrine (dKD) were used. Data are shown by box plots (n = 10 plants). The different letters indicate significant differences by Tukey’s HSD test (P < 0.03). Asterisks indicate significant differences from the value of FL with estradiol (+ER) (*t*-test, P < 0.01). (B) The relative number of rhizoids in (A). The average number of rhizoids in estradiol-free media (ER −) in each genotype was used for the normalization. Data are shown by box plots (n = 10 plants). The different letters indicate significant differences by Tukey’s HSD test (P < 0.03). Asterisks indicate significant differences from the value of FL with estradiol (+ER) (*t*-test, P < 0.01). (C) The number of rhizoids in the gemmae grown for 14 days in the medium supplemented with 1 µM estradiol. The gemmae of the inducible lines of full length MpNEK1-Citrine (FL), MpNEK1^K37E^-Citrine (K37E), and MpNEK1^dKD^-Citrine (dKD) were used. Data are shown by box plots (n indicates the number of plants). The different letters indicate significant differences by Tukey’s HSD test (P < 0.03).

To investigate the accumulation and subcellular localization of kinase-deficient MpNEK1, the inducible lines were grown in the medium with or without 1 µM estradiol and observed under confocal microscopy (Fig. 12). Because the inducible lines of kinase-deficient MpNEK1 develop larger thalli and a lot of rhizoids than the full length inducible lines, they provide good opportunity to analyze localization of MpNEK1 in the growing organs and more healthy cells. Estradiol treatment induced accumulation of MpNEK1^K37E^ and MpNEK1^dKD^ in the granules throughout the thallus as the full length MpNEK1 (Fig. 12A).

**Fig. 12.**
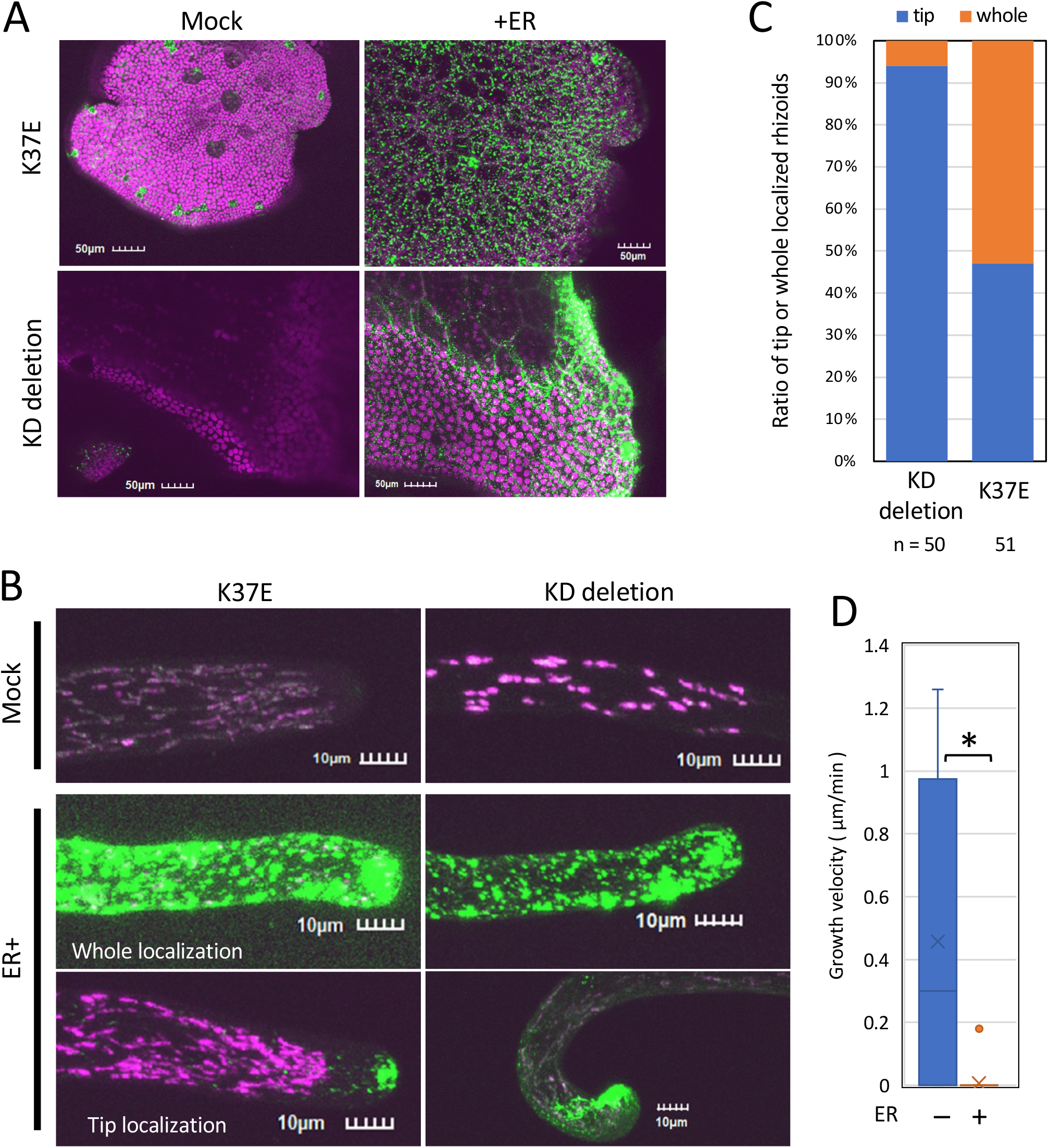
Localization of kinase-deficient MpNEK1-Citrine (A) The gemmae of the inducible lines of MpNEK1^K37E^-Citrine (K37E) or MpNEK1^dKD^-Citrine (KD deletion) were planted in the medium with (+ER) or without 1 µM estradiol (Mock), grown for 4 days, and observed under a confocal microscope. Green: MpNEK1-Citrine, magenta: plastid autofluorescence. (B) The gemmae of the inducible lines of MpNEK1^K37E^-Citrine (K37E) or MpNEK1^dKD^-Citrine (KD deletion) were grown as in (A) and rhizoids were observed under a confocal microscope. Green: MpNEK1-Citrine, magenta: plastid autofluorescence. (C) Ratio of rhizoids with tip-localized or whole localized MpNEK1^dKD^-Citrine (KD deletion) or MpNEK1^K37E^-Citrine (K37E) after 120 h of estradiol treatment. (D) Growth velocity of rhizoids in the MpNEK1^K37E^-Citrine inducible lines incubated for 48 hours with (+ER) or without 1 µM estradiol (-ER). Data are shown by box plots (n = 30 rhizoids). An asterisk indicates significant difference (*t*-test, P < 0.01).

We observed localization of MpNEK1^K37E^ and MpNEK1^dKD^ in rhizoids (Fig. 12B). Some rhizoids showed granular accumulation throughout rhizoids (whole localization), while others showed preferential localization at the apex of rhizoids (tip localization). We quantified the number of rhizoids with the whole localization or tip localization (Fig. 12C). Tip-localized rhizoids were approximately 47% in MpNEK1^K37E^ while 94% in MpNEK1^dKD^. Thus, MpNEK1^dKD^ exhibited relatively normal localization compared with MpNEK1^K37E^ in the inducible condition.

Next, we measured the growth velocity of rhizoids in the MpNEK1^K37E^ inducible lines (Fig. 12D). The rhizoid growth was almost stopped after 48 hours of estradiol treatment. This is comparable to the effect of the full length inducible lines. Therefore, excess accumulation of kinase-dead MpNEK1 also suppressed rhizoid elongation.

## Discussion

Here, we established and characterized induction system of MpNEK1 as a new tool to analyze plant NEK function. Estradiol-based XVE induction system has been widely used in angiosperms, but its application and characterization in *M. polymorpha* still remain limited. The efficient induction of Mp*NEK1* transcripts and overaccumulation of MpNEK1-Citrine fusion protein throughout plant body indicate that this system is very helpful for the functional analysis of MpNEK1 and other proteins in *M. polymorpha*.

We noticed several unexpected phenomena in which caution is required. MpNEK1-Citrine showed leaky expression in the meristem without estradiol. This might be due to the basal expression of artificial transcriptional activator XVE under the control of Mp*EF1α* promoter, that is highly active in meristematic tissues (Althoff, et al. 2014). Nonetheless, this leaky expression did not affect thallus growth, suggesting that effective threshold of MpNEK1. Since the leaky expression was varied in plants and transgenic lines, estrogen-free XVE-mediated transactivation might be stochastic and/or responsive to fluctuating physiological states.

Secondly, estrogen-resistant silenced plants emerged as survivors in the prolonged treatment especially at the lower concentrations of estradiol. The estrogen resistance could be transferred to the next generation gemmae, implying thallus-to-gemma movement of siRNA. Thus, it would be useful for the analysis of RNA silencing in *M. polymorpha*, while invokes concerns about treatment time and estradiol concentration to minimize the contamination of silenced plants.

MpNEK1 induction severely suppressed thallus growth probably through cell proliferation. There are several possible mechanisms of this growth-suppressive effect. Firstly, MpNEK1 may depolymerize microtubule-based mitotic apparatus including spindle and phragmoplast or affect cortical microtubule organization, which is required for directional growth prior to cell division. However, our microtubule imaging showed that overexpression of MpNEK1 did not severely affect mitotic apparatus and cortical microtubules. Secondly, overexpression of MpNEK1 may activate cell cycle checkpoints to prevent cell cycle progression. According to the experiment of EdU labeling, the cells through the S phase were reduced under the MpNEK1 overexpression. Furthermore, MpNEK1 induction reduced mitotic cells in the M phase. Thus, MpNEK1 induction may stop cell cycle in the G1 phase and eventually reduce the number of dividing cells. The *nimA* mutants of a filamentous fungus *Aspergillus nidulans* exhibit cell cycle arrest just before M phase, and conversely, overexpression of *NimA* causes premature chromosome condensation and aberrant spindle formation, resulting in cell cycle arrest (Oakley and Morris, 1983; Osmani, et al. 1988). Taking into accounts of remarkable promoter activity of Mp*NEK1* in the meristem (Otani, et al. 2018), it would be likely that MpNEK1 controls cell cycle progression and checkpoints.

Overexpression of MpNEK1 suppressed tip growth in rhizoids. Furthermore, overexpression altered localization of MpNEK1-Citrine from the rhizoid apex to the entire region as the time of estradiol treatment increased. This may reflect the overflow of MpNEK1 from the transport and localization machineries. As the consequence, tip growth was stopped in rhizoids with this dispersed localization pattern. Over-accumulated MpNEK1 may affect microtubule organization and/or microtubule-dependent processes during rhizoid growth. Because the longitudinal microtubules promote rhizoid elongation through organelle transport (Kanda, et al. 2022), it is plausible that MpNEK1 accumulation throughout rhizoids suppresses transport system (see the last paragraph).

Unexpectedly, we found growth suppression of thalli and rhizoids by kinase-deficient MpNEK1. This result demonstrates that growth suppression is independent of protein phosphorylation by MpNEK1. Therefore, the induction system is not suitable for the identification of target proteins phosphorylated by MpNEK1. Nevertheless, kinase-dead MpNEK1 is slightly milder than the wild type MpNEK1 in the estradiol induction system. The kinase activity of MpNEK1 is not essential for the growth suppression, but may have a minor contribution. Excessively accumulated MpNEK1 might trap interacting proteins and substrates in the granules to suppress thallus growth and rhizoid elongation in a kinase-independent manner. Animal NEK7 licenses the assembly and activation of NLRP3 inflammasome, a large protein complex for the activation of inflammatory caspases, independently of its kinase activity (He et al. 2016, Sharif et al. 2019). Furthermore, NEK7 is required for oligomerization of NLRP3 and its adaptor protein ASC, which form the granular structure in the cytosol (He et al. 2016, Sharif et al. 2019). Plant NEK may also function as the assembly factor of protein supercomplex and cytosolic granular structures.

In summary, MpNEK1 induction system is useful for the analyses of its function in cell proliferation. Furthermore, it would be helpful to visualize directional protein transport in rhizoids because we observed MpNEK1 particles moved to the acropetal and basipetal direction (Video S3-S5). The velocity of MpNEK1 particles is quite similar to that of armadillo-repeat kinesin (Kanda et al. 2022), about half of velocity of growing microtubule plus ends, suggesting this motility is not due to plus-end tracking but rather to the kinesin-dependent transport along microtubules. MpNEK1 overexpression has the potent growth effect than that of *Arabidopsis* NEK6, whose overexpression caused slight reduction of root growth (Takatani, et al. 2017). Comparing with MpNEK1, NEK6-Citrine fusion protein is not highly accumulated in the XVE system (H. Mo., unpublished result), implying the differential degradation efficiency of NEK proteins and the potential advantage of protein induction and analyses in *M. polymorpha*.

## Materials and Methods

### Plant materials and growth conditions

1. *M. polymorpha* accessions Takaragaike-1 (Tak-1, male) and Takaragaike-2 (Tak-2, female) were used as the wild type strains. Plants were grown on the half-strength Gamborg’s B5 medium solidified with 1% agar at 22°C under the light cycles with 16 hours of light and 8 hours of darkness.

### Plasmid construction and transformation

Plasmids were mainly constructed based on the Gateway technology (Life technologies). Primers used in the plasmid construction are listed in Table S1. To generate a plasmid of Mp*NEK1* induction by estradiol, the full-length MpNEK1 coding sequence in the pENTR/D-TOPO entry vector (Otani et al. 2018) was transferred by LR reaction (LR clonase [ enzyme mix, Life Technologies) into the Gateway binary vector pMpGWB344 (MpEF1α pro:XVE >>LexA operator:Gateway cassette).

To generate a plasmid of estradiol-inducible MpNEK1-mCitrine, *mCitrine* coding sequence was amplified by KOD One (Toyobo) and a primer set (Table S1) and was cloned into the *Asc*I site of the pENTR/D-TOPO entry vector containing full length MpNEK1 CDS (Otani et al., 2018) by In-fusion system (Takara). The resulting vector pENTR/D-TOPO-MpNEK1-mCitrine was subjected to LR reaction to transfer MpNEK1-Citrine fusion into the Gateway binary vector pMpGWB144 (MpEF1α pro:XVE >>LexA operator:Gateway cassette).

To generate a plasmid of estradiol-inducible kinase-deficient MpNEK1, pENTR/D-TOPO-MpNEK1-mCitrine was subjected to the inverse PCR-based mutagenesis using KOD mutagenesis kit (Toyobo) and primer sets (Table S1). The resulting vectors, pENTR/D-TOPO-MpNEK1^K37E^-mCitrine and pENTR/D-TOPO-MpNEK1^dKD^-mCitrine were subjected to LR reaction to transfer mutated MpNEK1-Citrine fusion into the Gateway binary vector pMpGWB144 (MpEF1α pro:XVE >>LexA operator:Gateway cassette).

The constructs, pMpGWB344-MpNEK1 (MpEF1α pro:XVE>>LexA operator:MpNEK1), pMpGWB144-MpNEK1-Citrine (MpEF1α pro:XVE>>LexA operator:MpNEK1-Citrine), pMpGWB144-MpNEK1^K37E^-Citrine (MpEF1α pro:XVE>>LexA operator:MpNEK1^K37E^-Citrine), and pMpGWB144-MpNEK1^dKD^-Citrine (MpEF1α pro:XVE>>LexA operator:MpNEK1^dKD^-Citrine) were transformed into F1 sporelings derived from sexual crosses between Tak-2 and Tak-1 by *Agrobacterium*-mediated method (Ishizaki et al., 2008). The pMpGWB344-MpNEK1 construct was transformed into the regenerating thalli of CaMV35Spro:Citrine-MpTUB2 plants harboring pMpGWB105-MpTUB2 construct (Otani et al. 2018) according to the method of Kubota et al., 2013. Chlorsulfuron and hygromycin was used for the selection of transformants in pMpGWB344 and pMpGWB144, respectively. In pMpGWB344 and pMpGWB144, the artificial transcriptional activator XVE is expressed under the control of Mp*EF1*_α_ promoter. XVE translocates to the nucleus in the presence of estradiol and binds to the LexA operator to induce the expression of MpNEK1 and MpNEK1-Citrine, respectively.

### RT-qPCR

Total RNA was isolated from the thalli of the wild type and transgenic plants. For each sample, 0.5 µg of total RNA was reverse transcribed to cDNA using ReverTra Ace reverse transcriptase (Toyobo) according to the manufacturer’s protocol. Real-time PCR was performed on a thermal cycler Dice Real Time System (Takara) using KOD SYBR qPCR Kit (Toyobo) according to the manufacturer’s protocol. Transcript levels of Mp*EF1*_α_ and Mp*ACT* were used as a reference for normalization (Saint-Marcoux et al., 2015). Primers used in RT-qPCR are listed in Table S1. RT-qPCR experiments were performed using four biological replicates.

### Microscopy

The morphology of gemmalings and rhizoids was observed under a stereo microscope S8APO0 (Leica Microsystems) or a light microscope DM5000B (Leica) equipped with a CCD camera

DFC500 (Leica). To analyze the localization of MpNEK1-Citrine, plants were grown on half-strength B5 medium for 3 and 5 days under the cycles with 16 hours of light and 8 hours of darkness. For live imaging, half-strength B5 agar medium was solidified in the glass bottom dish (35 mm in diameter x 10 mm in thickness, Thickness No. 1, Matsunami) and then the central region of agar medium on the bottom glass strip was removed by tweezers. This region was poured with about 200 µL of melted half-strength B5 agar medium to generate a thin solidified medium. The gemmae were planted and grown in the central thin medium for 2-7 days. The details of this method will be described elsewhere by H.M. Gemmalings were observed under a FV1200 confocal laser-scanning microscope (Olympus) equipped with a high-sensitivity GaAsP detector and silicone oil objective lenses (30×, 60× Olympus) or FV3000 (Olympus) equipped with high-sensitivity GaAsP detectors and oil immersion objective lenses (40x, NA = 1.4; 60x, NA = 1.42, Olympus). Silicone oil (SIL300CS, Olympus) or immersion oil (F30CC, Olympus) was used as immersion media for the objective lenses. The samples were excited at 473 nm and 559 nm (laser diode) and the emission was separated using a FV12-MHSY SDM560 filter (490-540 nm, 575-675 nm, Olympus) in FV1200. The images were analyzed using ImageJ (National Institutes of Health, USA). For estradiol treatment, an aliquot (200[300 µL) of 1/2 B5 liquid medium supplemented with estradiol at a concentration of 1 µM or 10 µM was added to the gemmalings for the induction of Mp*NEK1*. As a control, same amount of 1/2 B5 liquid medium supplemented with same concentration of solvent (DMSO, less than 0.1%) was applied.

### EdU staining

S-phase cells were visualized using Click-iT EdU Imaging Kits (Life Technologies) according to the manufacturer’s instruction. Gemmae were incubated on the half-strength Gamborg’s B5 agar medium supplemented with or without 1 µM β-estradiol for 4 days. Alternatively, for 1-day treatment, gemmae were incubated on the agar medium without 1 µM β-estradiol for 3 days and then transferred to and incubated in the agar medium with 1 µM β-estradiol for 1 day. Gemmalings were then transferred to a solution containing 10 µM 5-ethynyl-2-deoxyuridine (EdU) and incubated for 1 hour. Plants were fixed with 3.7 % formaldehyde solution in phosphate-buffered saline (PBS) for 20 min. EdU incorporated into DNA was labelled with Alexa Fluor 488-azide-containing Click-iT reaction cocktail in the dark for 30 min. EdU-labelled cells were observed using a confocal laser scanning microscope, FV1200 (Olympus). The maximum Z-projection images were created using the ImageJ software (Schneider et al. 2012).

## Abbreviations

EdU: 5-ethynyl-2-deoxyuridine
EF1α: elongation factor1α
NEK: NIMA-related kinase
NIMA: never in mitosis A WDL, WAVE DANPENED-LIKE, XVE: LexA-VP16-ER artificial transcription activator
WT: wild type

## Acknowledgements

We thank Drs. Ryuichi Nishihama and Kimitsune Ishizaki for their advice on EdU staining and transformation.

## Author contributions

HMa and HMo designed the research; HMa performed experiments and analyzed the data; YY and TK developed XVE system; HMa and HMo wrote the manuscript; HMa, YY, TK, TT and HMo revised the manuscript.

## Data availability

The data of this study are available upon request from the corresponding author, Hiroyasu Motose.

## Supplementary data

Fig S1. Effects of estradiol on the accumulation of Mp*NEK1* transcripts in the wild type and transgenic lines introduced with pMpGWB344-Mp*NEK1* (*XVE::*Mp*NEK1*).

The 10-day old plants of the wild type (WT) and the transgenic lines were incubated for 1 day in the medium with (+) or without 10 µM estradiol (-). Total RNA was extracted and subjected to RT-qPCR. All transcript levels are relative to that of mock-treated wild type plants. Mp*EF1*_α_ and Mp*ACT* were used as the control genes. Columns and error bars indicate mean values and standard errors, respectively (n indicates the number of biological replicates).

Fig S2. Effect of estradiol on thallus growth of the wild type.

(A) The gemmae of the wild type (Tak-1) were planted in the agar medium with (+ER) or without 10 µM estradiol (Mock) and grown for 3, 11, and 20 days.

(B) Time course of growth of the wild type with (+ER) or without 10 µM estradiol (Mock).

(C) Quantification of thallus growth of the wild type with (+ER) or without 10 µM estradiol (Mock). The mean projection area of thalli (n = 10 plants) was quantified by ImageJ. Circles and error bars indicate mean values and standard deviations, respectively.

Fig S3. Effect of estradiol on thallus growth of the transgenic line #6 with *XVE::*Mp*NEK1*.

(A) The gemmae of the line (#6) were planted in the agar medium with (+ER) or without 10 µM estradiol (Mock) and grown for 3, 11, and 20 days.

(B) Time course of growth of the line (#6) with (+ER) or without 10 µM estradiol (Mock).

(C) Quantification of thallus growth of the line (#6) with (+ER) or without 10 µM estradiol (Mock).

The mean projection area of thalli (n = 10 plants) was quantified by ImageJ. Circles and error bars indicate mean values and standard deviations, respectively.

Fig S4. Effect of estradiol on the accumulation of Mp*NEK1* transcripts in the wild type and estradiol-resistant transgenic plants harboring *XVE::*Mp*NEK1*.

The 10-day old plants of the wild type (WT) and estradiol-resistant plants derived from line #8 and #2 were incubated for 1 day in the medium with (+) or without 10 µM estradiol (-). Total RNA was extracted and subjected to RT-qPCR. All transcript levels are relative to that of mock-treated wild type plants. Mp*EF1*_α_ were used as the control genes. Data are shown by box plots (n = 2 biological replicates).

Fig S5. Effect of estradiol on the growth of MpNEK1-Citrine inducible lines

The gammae of the wild type (WT) and the MpNEK1-Citrine inducible lines were planted in the agar medium supplemented with (+ER) or without 1 µM estradiol (Mock) and grown for 15 days.

Fig S6. Expression and localization of MpNEK1-Citrine in the meristem of thallus without estradiol.

Left panel shows a light field image merged with a confocal image in the right panel. Green; MpNEK1-Citrine, magenta; plastid autofluorescence.

Fig S7. Effect of estradiol on the growth of the wild type and MpNEK1-Citrine inducible lines

The gammae of the wild type (WT) and the MpNEK1-Citrine inducible lines were planted in the agar medium supplemented with (+ER) or without 1 µM estradiol (Mock) and grown for 7, 14, and 21 days.

Video S1. Localization of MpNEK1-Citrine in a gemmaling grown for 4 days in the medium supplemented with 1µM estradiol (same plant shown in Fig. 6B right panels).

Video S2. Localization and dynamics of MpNEK1-Citine in a rhizoid of MpNEK1-Citrine inducible line treated with estradiol for 24 hours.

MpNEK1-Citrine inducible line was grown without estradiol for 3 days and then applied with 1 µM estradiol and grown for 24 hours. Green: MpNEK1-Citrine, magenta: plastid autofluorescence (same rhizoid shown in Fig. 7C).

Video S3. Localization and dynamics of over-accumulated MpNEK1-Citine in a rhizoid of MpNEK1-Citrine inducible line treated with estradiol for 24 hours.

MpNEK1-Citrine inducible line was grown without estradiol for 3 days and then applied with 1 µM estradiol and grown for 24 hours. A rhizoid highly accumulating MpNEK1-Citrine was observed. Green: MpNEK1-Citrine, magenta: plastid autofluorescence.

Video S4. Localization and dynamics of MpNEK1-Citine in a rhizoid of MpNEK1-Citrine inducible line treated with estradiol for 48 hours.

MpNEK1-Citrine inducible line was grown without estradiol for 3 days and then applied with 1 µM estradiol and grown for 48 hours. Green: MpNEK1-Citrine, magenta: plastid autofluorescence (same rhizoid shown in Fig. 7C).

Video S5. Localization and dynamics of MpNEK1-Citine in a rhizoid of MpNEK1-Citrine inducible line treated with estradiol for 48 hours.

MpNEK1-Citrine inducible line was grown without estradiol for 3 days and then applied with 1 µM estradiol and grown for 48 hours. Green: MpNEK1-Citrine, magenta: plastid autofluorescence (same rhizoid shown in Fig. 8A).

Video S6. Localization and dynamics of MpNEK1-Citine in a rhizoid of MpNEK1-Citrine inducible line treated with estradiol for 120 hours.

MpNEK1-Citrine inducible line was grown without estradiol for 3 days and then applied with 1 µM estradiol and grown for 120 hours. Green: MpNEK1-Citrine, magenta: plastid autofluorescence (same rhizoid shown in Fig. 7C).

Video S7. Localization of MpNEK1-Citine in a rhizoid of MpNEK1-Citrine inducible line treated with estradiol for 120 hours.

MpNEK1-Citrine inducible line was grown without estradiol for 3 days and then applied with 1 µM estradiol and grown for 120 hours.Green: MpNEK1-Citrine, magenta: plastid autofluorescence.

