## Supplementary material for "Overexpression of NIMA-related kinase suppresses cell proliferation and tip growth in a liverwort *Marchantia polymorpha*": Suppl Fig

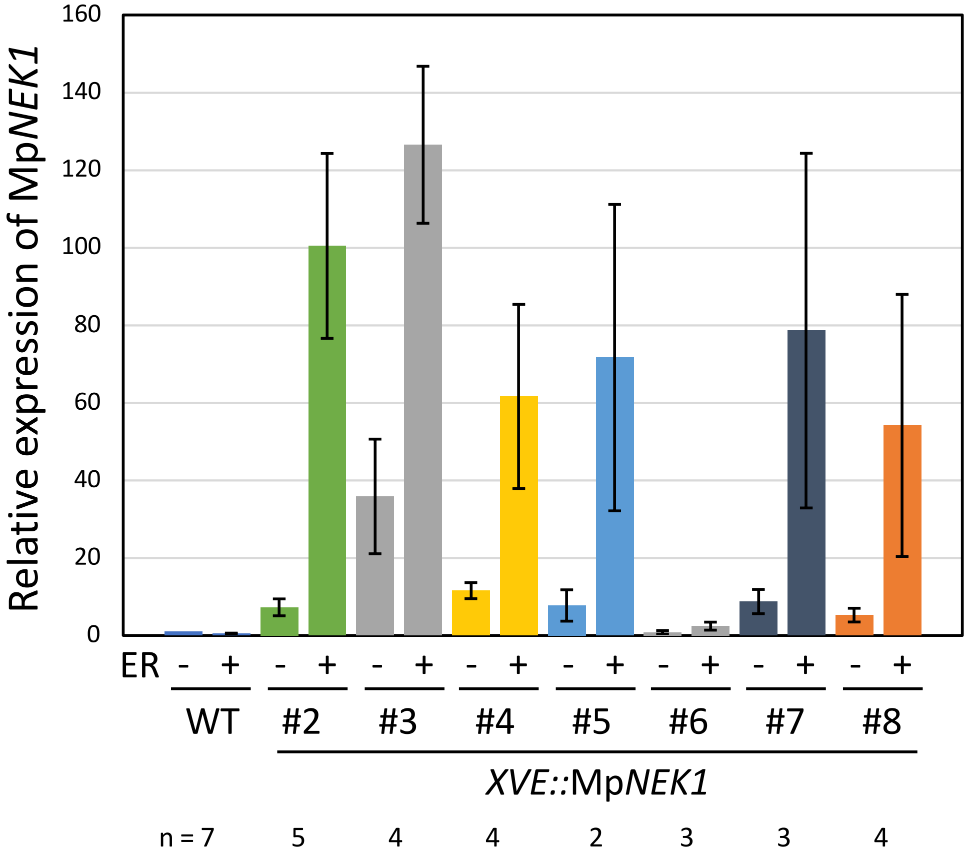


Fig S1. Effects of estradiol on the accumulation of Mp*NEK1* transcripts in the wild type and transgenic lines introduced with *XVE::*Mp*NEK1*.
The 10-day old plants of the wild type (WT) and the transgenic lines were incubated for 1 day in the medium with (+) or without 10 µM estradiol (-). Total RNA was extracted and subjected to RT-qPCR. All transcript levels are relative to that of mock-treated wild type plants. Mp*EF1α* and Mp*ACT* were used as the control genes. Columns and error bars indicate mean values and standard errors, respectively (n indicates the number of biological replicates).

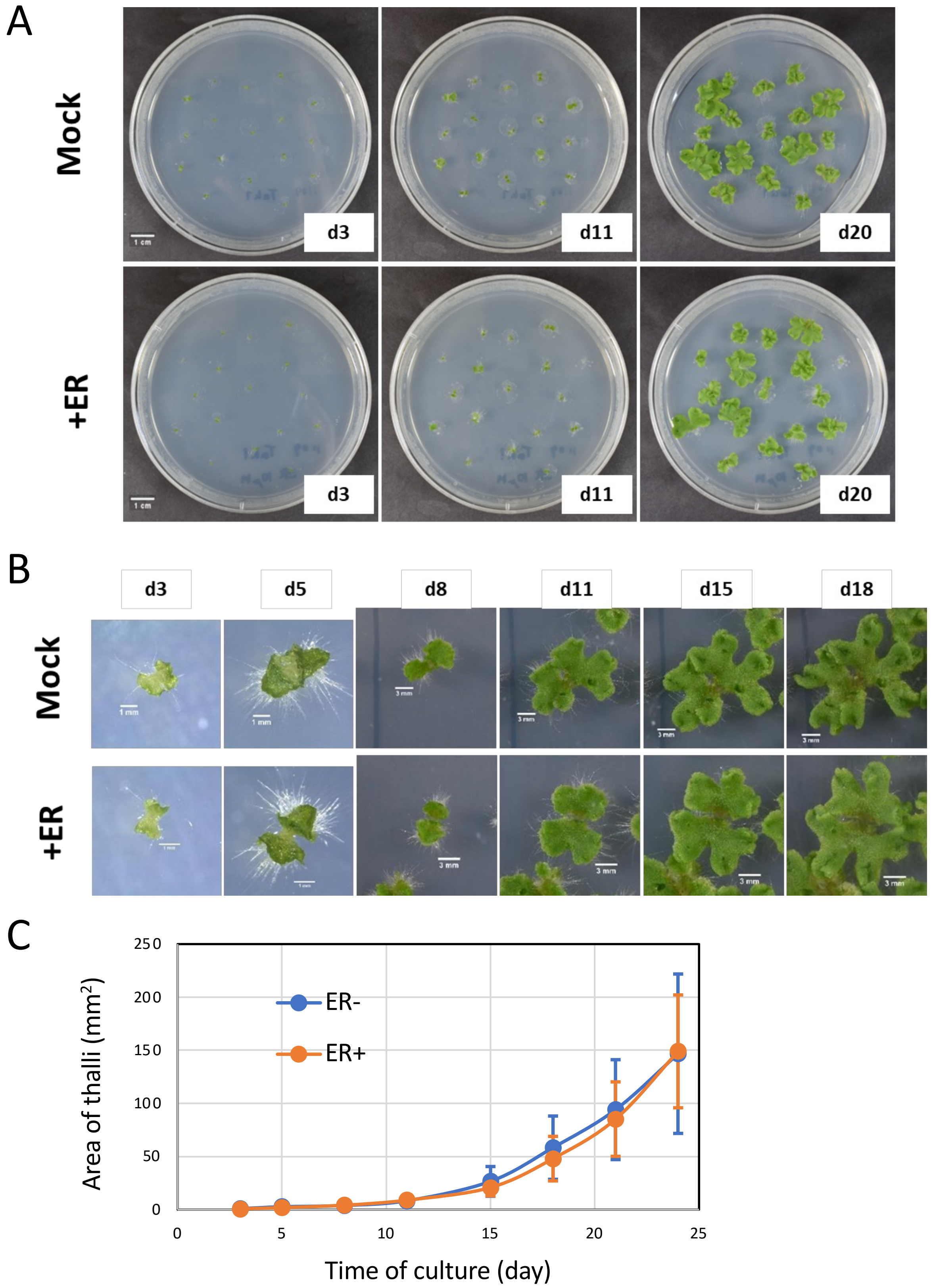


Fig S2. Effect of estradiol on thallus growth of the wild type.

(A) The gemmae of the wild type (Tak-1) were planted in the agar medium with (+ER) or without 10 µM estradiol (Mock) and grown for 3, 11, and 20 days.

(B) Time course of growth of the wild type with (+ER) or without 10 µM estradiol (Mock) .

(C) Quantification of thallus growth of the wild type with (+ER) or without 10 µM estradiol (Mock). The mean projection area of thalli (n = 10 plants) was quantified by ImageJ. Circles and error bars indicate mean values and standard deviations, respectively.

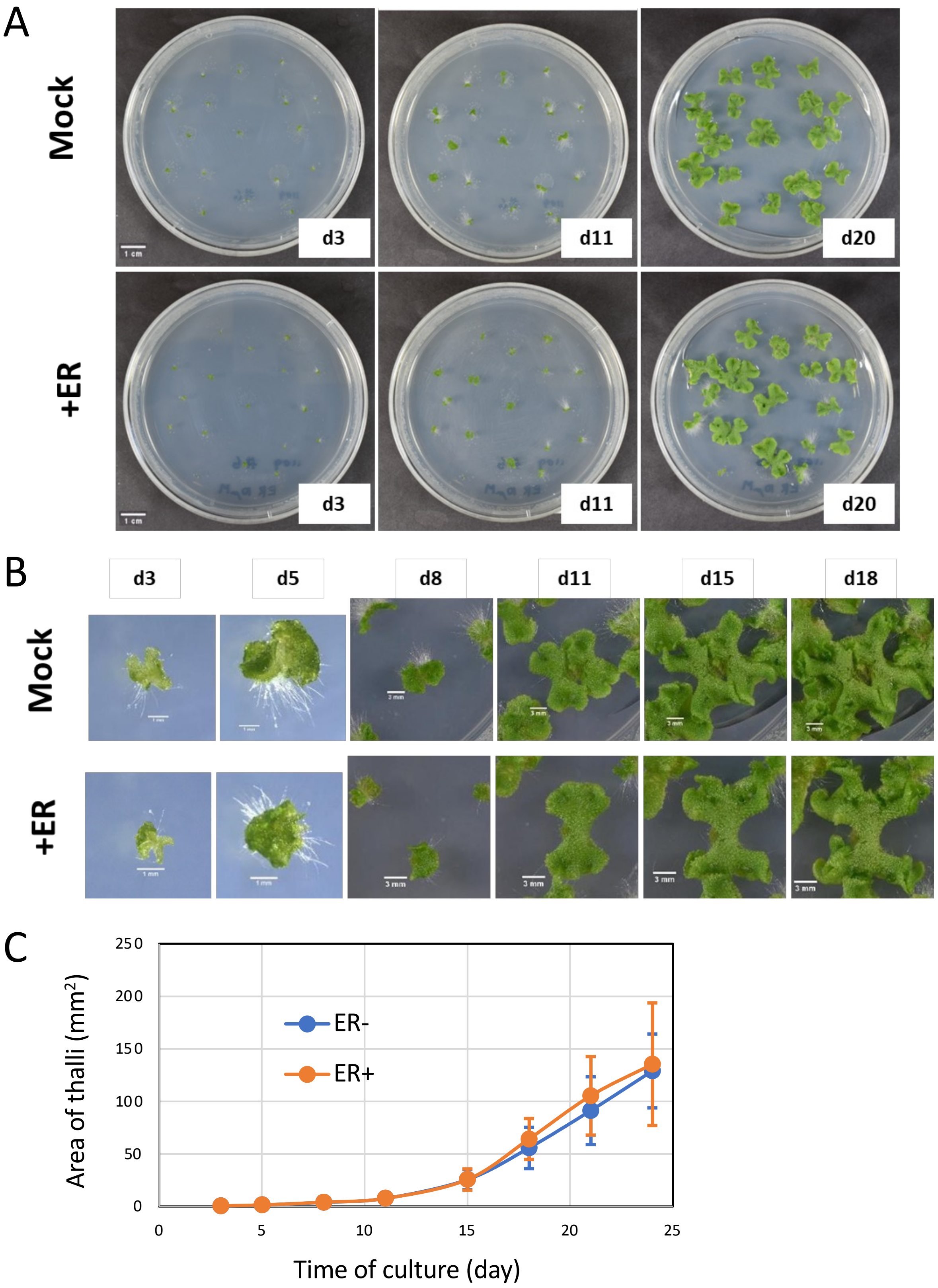


Fig S3. Effect of estradiol on thallus growth of the transgenic line #6 with *XVE::*Mp*NEK1*.

(A) The gemmae of the line (#6) were planted in the agar medium with (+ER) or without 10 μM estradiol (Mock) and grown for 3, 11, and 20 days.

(B) Time course of growth of the line (#6) with (+ER) or without 10 μM estradiol (Mock) .

(C) Quantification of thallus growth of the line (#6) with (+ER) or without 10 μM estradiol (Mock). The mean projection area of thalli (n = 10 plants) was quantified by ImageJ. Circles and error bars indicate mean values and standard deviations, respectively.

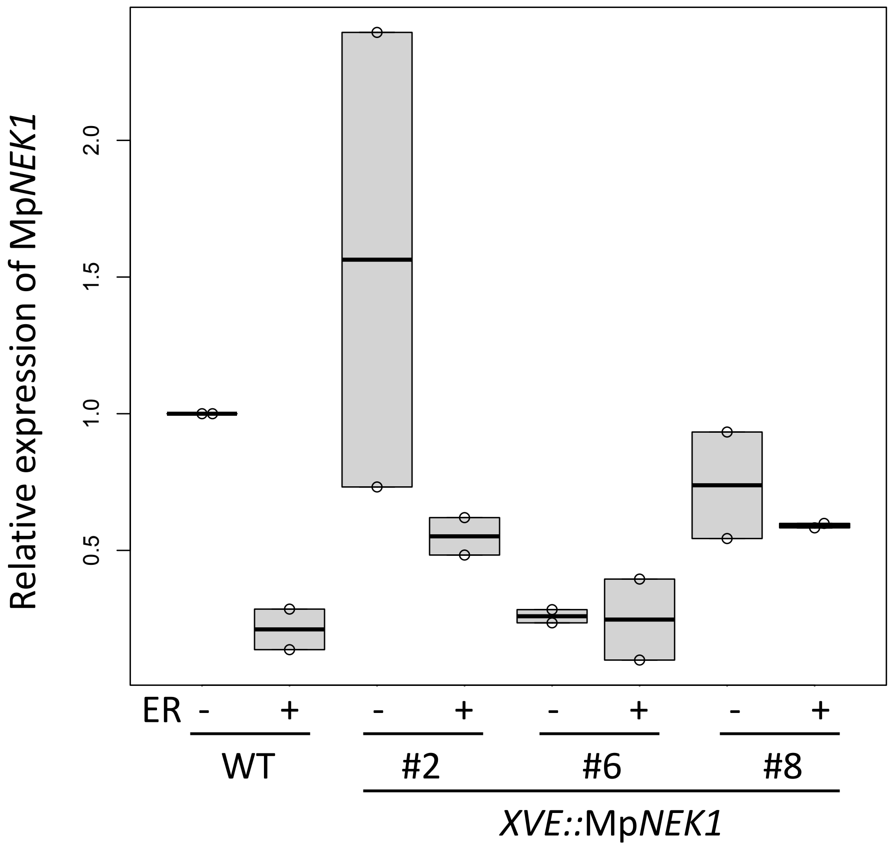


Fig S4. Effect of estradiol on the accumulation of Mp*NEK1* transcripts in the wild type and estradiol-resistant transgenic plants harboring *XVE::*Mp*NEK1*.

The 10-day old plants of the wild type (WT) and estradiol-resistant plants derived from line #8 and #2 were incubated for 1 day in the medium with (+) or without 10 µM estradiol (-). Total RNA was extracted and subjected to RT-qPCR. All transcript levels are relative to that of mock-treated wild type plants. Mp*EF1α* was used as the control genes. Data are shown by box plots (n = 2 biological replicates).

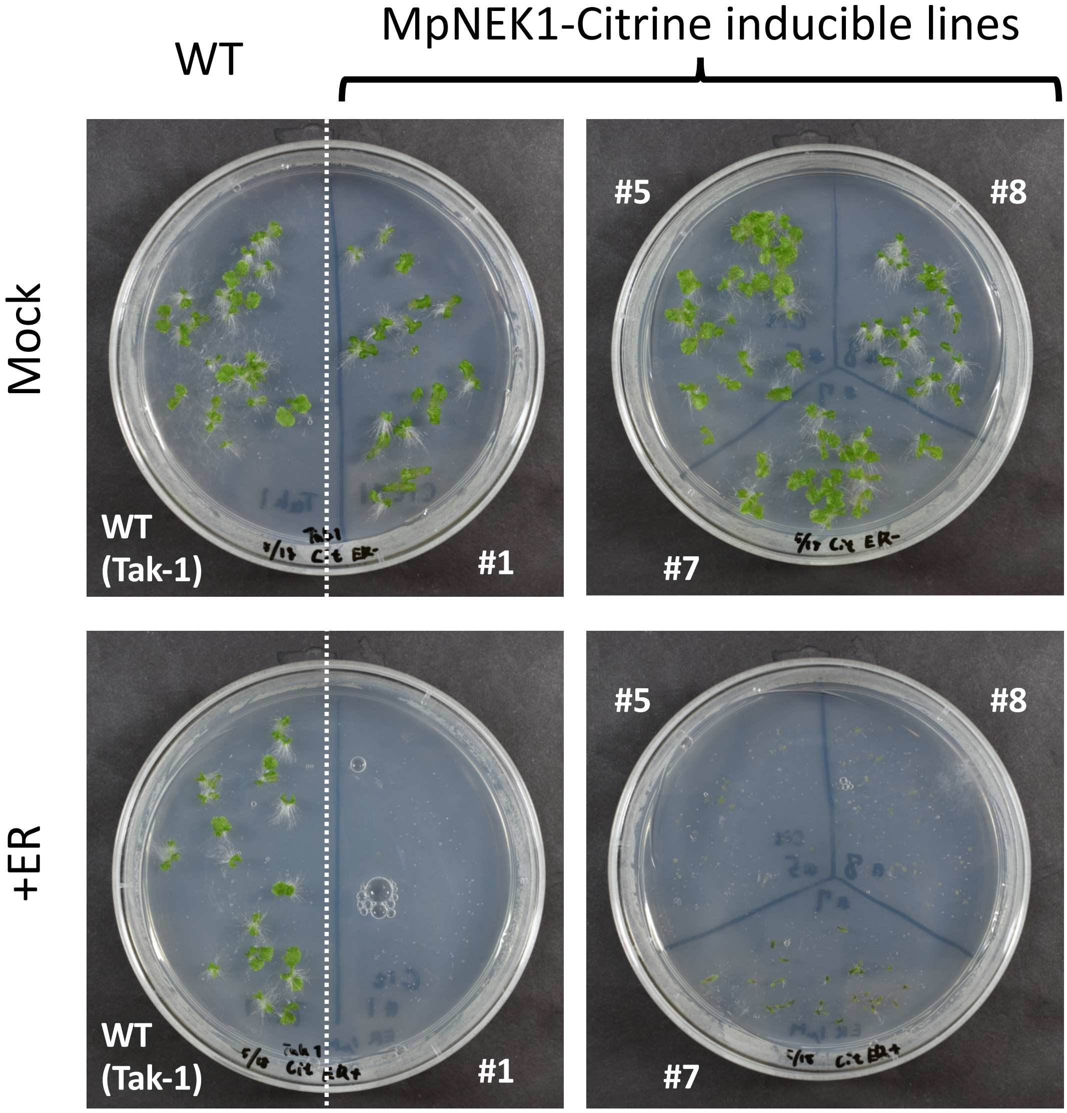


Fig S5. Effect of estradiol on the growth of MpNEK1-Citrine inducible lines

The gammae of the wild type (WT) and the MpNEK1-Citrine inducible lines were planted in the agar medium supplemented with (+ER) or without 1 μM estradiol (Mock) and grown for 15 days.

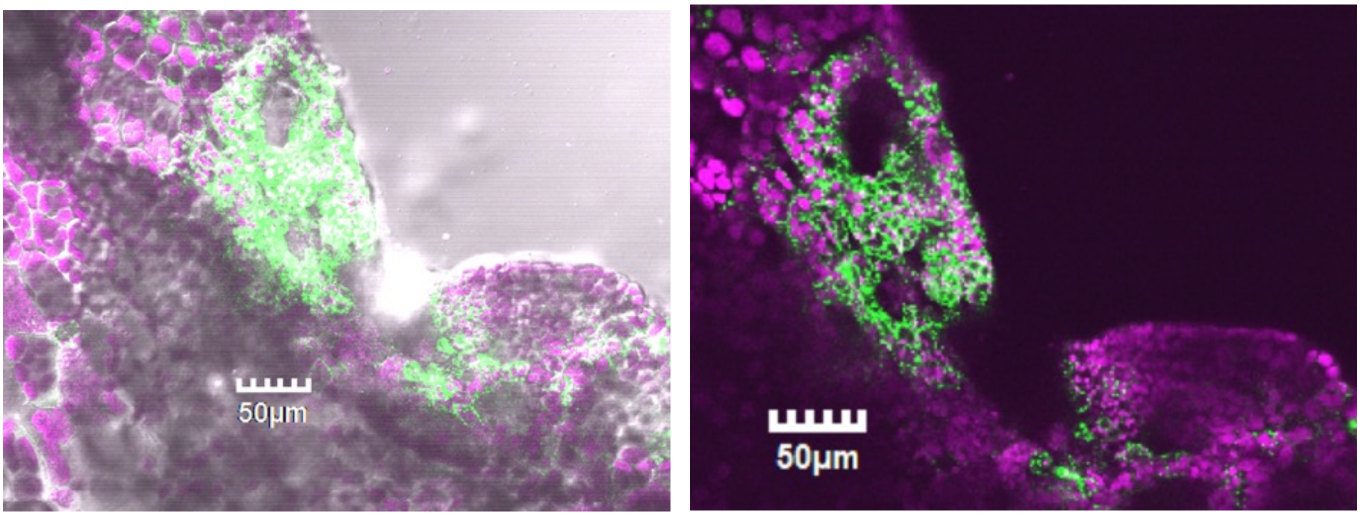


Fig S6. Expression and localization of MpNEK1-Citrine in the meristem of thallus without estradiol.

Left panel shows a light field image merged with a confocal image in the right panel. Green; MpNEK1-Citrine, magenta; plastid autofluorescence.

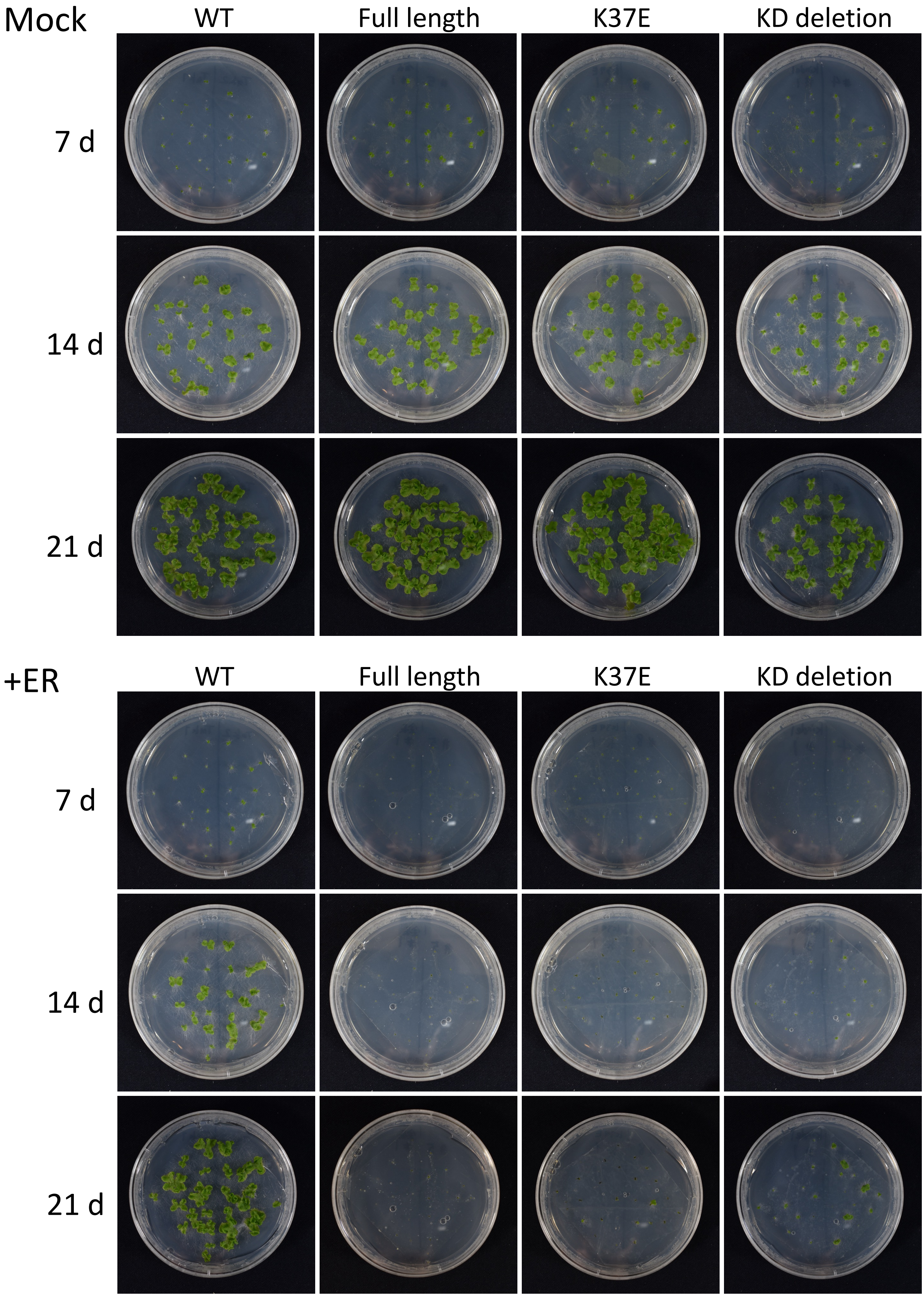


Fig S7. Effect of estradiol on the growth of the wild type and MpNEK1-Citrine inducible lines

The gammae of the wild type (WT) and the MpNEK1-Citrine inducible lines were planted in the agar medium supplemented with (+ER) or without 1 μM estradiol (Mock) and grown for 7, 14, and 21 days.
