## Supplementary material for "Overexpression of NIMA-related kinase suppresses cell proliferation and tip growth in a liverwort *Marchantia polymorpha*": Suppl Table

Table S1. Primers used in this study

| Primer | Sequence | Purpose |
|---|---|---|
| MpNEK-F1-RT(50-69) | GCGCCTTTGGATCTGCAATA | RT-qPCR |
| MpNEK-R1-RT(205-224) | ACCCAGGCTTCCTTGTAGTC | RT-qPCR, colony PCR & sequencing of XVE vectors |
| MpNEK-F2-RT-rev | TTCCAGCAGAAcGACGACTG | RT-qPCR, colony PCR & sequencing of XVE vectors |
| MpNEK-R2-RT(2144-63) | GTAGCTCAGGCTGTCCTTGG | RT-qPCR |
| MpEF1-F1 | AAGCCGTCGAAAAGAAGGAG | RT-qPCR |
| MpEF1-R1 | TTCAGGATCGTCCGTTATCC | RT-qPCR |
| MpACT-F1 | AGGCATCTGGTATCCACGAG | RT-qPCR |
| MpACT-R1 | ACATGGTCGTTCCTCCAGAC | RT-qPCR |
| XVE-after35S-F | ACTCTAGCCTCGAGGCGCGC | Colony PCR & sequencing of XVE vectors |
| MpNEK-F5 | ATGCAGCCGGACTACGACGAG | Colony PCR & sequencing of XVE vectors |
| MpNEK-F6 | TGGACCTGCCCGGCAAGATGG | Colony PCR & sequencing of XVE vectors |
| Cit-F-InF-AscI | agtaagggtgggcgcgccgacATGGTGAGCAAGGGCGAGGAGCT | In-Fusion cloning of Citrine |
| Cit-R-InF-AscI | agctgggtcggcgcgTTACTTGTACAGCTCGTCCATGCCG | In-Fusion cloning of Citrine |
| MpNEK1-F-K37E | gAAAAGATCCGTCTCGCTCGTCAGACG | Innverse PCR-based mutagenesis (K37E substitution) |
| MpNEK1-R-L36 | GAGGACATACTTCTTCTTTTCGAGCT | Innverse PCR-based mutagenesis (K37E substitution) |
| MpNEK1-F-Ctail+start | atgCAACCGTATATCACTCAGTGCCGGTT | Innverse PCR-based mutagenesis (deletion of kinase domain) |
| ENTR-D-R(+start) | CATGGTGAAGGGGGCGGCCG | Innverse PCR-based mutagenesis (deletion of kinase domain) |
| Citrine-R-176 | GAAGGTGGTCACGAGGGTGG | Colony PCR & sequencing of XVE vectors |
